# USP9X is a mechanosensitive deubiquitinase that controls tumor cell invasiveness and drug response through YAP stabilization

**DOI:** 10.1101/2024.08.29.610314

**Authors:** Pierric Biber, Alexandrine Carminati, Christophe A. Girard, Walaa Mohager, Mickael Ohanna, Margaux Lecacheur, Océane Bouvet, Stéphane Audebert, Mehdi Khaled, Sophie Tartare-Deckert, Marcel Deckert

**Author notes:** Equal contribution (Pierric Biber, Alexandrine Carminati). **Corresponding Author:** Marcel Deckert, Inserm UMR1065/C3M, 151 Route de Ginestière BP2 3194, F-06204 Nice cedex 3.

## Abstract

Post-translational modification by ubiquitin is crucial for protein turnover. Deubiquitinases (DUBs) remove ubiquitin chains from target proteins to prevent their degradation by the proteasome, thus acting as gatekeepers of protein homeostasis alongside the ubiquitin-proteasome system (UPS). Tumor cells exhibit remarkable plasticity, enabling them to adapt to anticancer treatments and the conditions of the tumor microenvironment, including mechanical cues from the extracellular matrix (ECM). However, the role of DUBs in mechanotransduction remains unexplored. To identify DUBs involved in cancer cell mechanosignaling, we used melanoma cells grown on collagen matrices with varying stiffnesses and an activity-based ubiquitin probe to profile DUB activities. Our approach, combined with quantitative proteomics, revealed that ubiquitin-specific protease 9X (USP9X) is sensitive to ECM stiffness through discoidin domain receptors (DDR)/actomyosin signaling pathway. In silico analysis further indicated that the mechanosensor YAP is part of the USP9X interactome, and USP9X expression correlates with the YAP transcriptional signature in melanoma. We hypothesized that mechanical signals regulate YAP levels through USP9X DUB activity. Consistently, low collagen stiffness reduced YAP expression, and siRNA-mediated depletion or pharmacological inhibition of USP9X decreased YAP protein expression in tumor cells. Conversely, knockdown of the ubiquitin E3 ligase βTrCP increased YAP protein levels. Affinity purification of polyubiquitinated proteins using Tandem Ubiquitin Binding Entities (TUBEs) showed that combined USP9X and proteasome inhibition increased YAP poly-ubiquitination, revealing that USP9X deubiquitinates YAP to prevent its proteasomal degradation. Targeting USP9X impaired stiffness-mediated responses, including YAP nuclear translocation and transcriptional activity, cell migration and invasion, and drug resistance. An experimental metastasis assay showed that stable knockdown of USP9X impaired melanoma cell lung colonization. Finally, targeting USP9X in a syngeneic BRAF-mutant melanoma model counteracted targeted therapy-induced ECM remodeling, enhanced treatment efficacy, and delayed tumor relapse. Our findings reveal a novel role of USP9X in cancer cell mechanobiology and drug resistance through stiffness-dependent stabilization of the oncoprotein YAP, proposing USP9X as a targetable “mechano-DUB” in cancer.

## Introduction

In cancer cells, genetic alterations, oncogenic pathways, and high proliferation rates lead to increased protein synthesis and addiction to mechanisms that maintain protein homeostasis, such as the ubiquitin-proteasome system (UPS) (1–5). Ubiquitination, a post-translational modification of proteins, constitutes a complex signaling system that coordinates essential cellular functions and maintains protein homeostasis (6). While mono-ubiquitination primarily modifies protein activity and regulates intracellular transport, K48-linked polyubiquitination is the principal mechanism governing the degradation of cellular proteins via the UPS. This system comprises a proteolytic complex (the 26S proteasome) and a cascade of enzymes (E1, E2, and E3) that activate ubiquitin residues and attach them to target proteins (7). Opposing this ubiquitin-conjugation process by E3 ligases, deubiquitination is carried out by deubiquitinating enzymes (DUBs), a group of ubiquitin-specific proteases that can remove one or more ubiquitin molecules from target proteins or disassemble entire polyubiquitin chains, thereby preventing their degradation by the proteasome (8,9). Aberrant expression or activity of DUBs has been implicated in pathological conditions, including cancer, making them promising targets for the development of novel molecular therapies (10,11). The accumulation of numerous oncoproteins related to tumor cell survival, plasticity and invasion, is regulated post-translationally by the UPS (12–14). Increased deubiquitination of proteasome substrates by DUBs protects these protumorigenic factors from degradation and promotes their accumulation leading to tumor growth, dissemination and therapeutic resistance (8,11,15).

Tumors are characterized by abnormal extracellular matrix (ECM) deposition and greater stiffness than healthy tissue that generates mechanical stresses, which correlates with tumor progression and aggressiveness (16–18). Tumor rigidity is sensed by cells through dedicated mecanosensing pathways, including downstream effectors of the Hippo pathway, Yes-associated protein (YAP) and its homolog TAZ. YAP/TAZ are key regulators of mechanotransduction, translating extracellular mechanical forces into transcriptional responses (19,20). Tumor cells exhibit remarkable plasticity, enabling them to adapt to anticancer treatments and the conditions of the tumor microenvironment, including mechanical cues from stiff ECM (21,22). Cutaneous melanoma is the skin cancer with the highest mortality rate, characterized by its high metastatic potential and resistance to treatment. Genetic alterations in the *BRAF*, *NRAS*, or *NF1* genes define melanoma subtypes and lead to constitutive activation of the MAPK pathway in over 75% of cases (23). Therapies targeting mutant BRAF and MEK and immunotherapies have led to improved survival in patients with metastatic disease (24–26). However, most patients do not respond to therapies or relapsed and require additional treatments. Activating mutations in the MAPK pathway induce melanocyte transformation, leading to primary tumors characterized by high mutational loads and extensive intratumoral phenotypic heterogeneity (27–29). Melanoma cell phenotypic plasticity and non-genetic heterogeneity actively contribute to drug resistance and relapse through transcriptional and epigenetic reprogramming in response to environmental challenges or therapeutic stress (21,30–32). Recent findings reveal that interactions between cancer cells and the ECM contribute to adaptive and acquired resistance to targeted therapy by creating a drug-protective niche for melanoma cells (33–36). MAPK pathway inhibition triggers collagen and fibronectin production (37–39) and ECM remodeling by melanoma cells lead to a cross-linked collagen matrix and tumor stiffening that triggers a feedforward loop dependent on the mechanotransducer YAP, resulting in therapeutic escape (35,40,41).

Several DUBs have been linked to melanoma progression and response to therapy (42), including USP13 that regulates MITF expression and melanoma growth (43), USP14, one of the three proteasome-associated DUBs that controls melanoma cell survival (44), and USP9X as an important regulator of ETS1 and NRAS expression (45). Thus, DUBs represent essential gatekeepers of protein homeostasis in cancer cells, including melanoma cells. However, whether they contribute to ECM stiffness-mediated tumor mechanical responses remains largely unexplored.

In this study, we used melanoma cells cultivated on collagen matrices with different stiffness, combined with an activity-based ubiquitin probe for profiling DUB activity and quantitative proteomics to identify USP9X as a DUB whose activity is modulated by ECM stiffness and actomyosin cytoskeleton remodeling. We also demonstrated that the mechanotransducer YAP is a substrate of the deubiquitinating activity of USP9X and that targeting USP9X mirrors the function of YAP in stiffness-induced melanoma cell migration, invasion, and therapeutic resistance *in vitro* and *in vivo*.

## Results

### ECM stiffness controls USP9X activity through a DDR/actomyosin pathway

To identify deubiquitinases whose activity is modulated by ECM rigidity, we first cultured a melanoma line (1205Lu) and melanoma short-term cultures (MM029 and MM099) on soft or stiff collagen substrates. Extracted proteins from these cultures were labeled using an activity-based HA-tagged ubiquitin probe that covalently binds DUB active site (Fig. 1A). This method works by using a probe formed by a vinylsulfone moiety fused to a ubiquitin and a hemagglutinin (HA) tag. The vinylsulfone moiety coupled to the ubiquitin, allows the recognition of the probe by active DUBs in the lysate to which it will covalently bind (44). Collagen stiffness impacted the global activation pattern of DUBs in melanoma cells (Fig. 1B). To extend our observations, we applied the same protocol to cell lines from other solid cancers: lung (A549), breast (MDA-MB-231), and pancreas (PANC1). For these different lines, we made the same observation, namely a difference in the overall pattern of DUBs activity as a function of stiffness (Supplementary Fig. 1). Given that the transmission of mechanical signals within the cell is dependent on the contraction of the actomyosin cytoskeleton, we used the myosin II inhibitor Blebbistatin. We observed that Blebbistatin treatment reduces the global activation pattern of DUBs in cells grown on a rigid matrix, compared to soft conditions (Fig. 1C). To individually identify DUBs whose activity is impacted by ECM stiffness we again applied this protocol to immunoprecipitate HA-Ub labelled DUBs on lysates from 1205Lu melanoma cells plated on soft and stiff collagen substrates. Using quantitative proteomics, we identified several DUBs with increased activity either in soft or stiff conditions (Fig. 1D, Supplementary Table 1). Among these DUBs, the activity of USP9X was increased by collagen stiffening (Fig. 1E). Conversely, stiffness-induced USP9X activity was blocked by Blebbistatin (Fig. 1F). The collagen discoidin domain receptor 1 (DDR1) has been involved in mechanical signaling (46) and we recently revealed the role of DDR1 and DDR2 in ECM-mediated drug resistance in melanoma cells (36). We thus investigated whether DDR1 and DDR2 (DDR) targeting affects ECM stiffness-mediated USP9X activity. We found that pharmacological or genetic inhibition of DDR markedly reduces USP9X activation on stiff collagen compared to what is observed on soft collagen (Fig. 1G). Together, the data indicate that the DUB activity of USP9X is increased on stiff collagen matrices through a DDR/actomyosin pathway.

**Figure 1.**
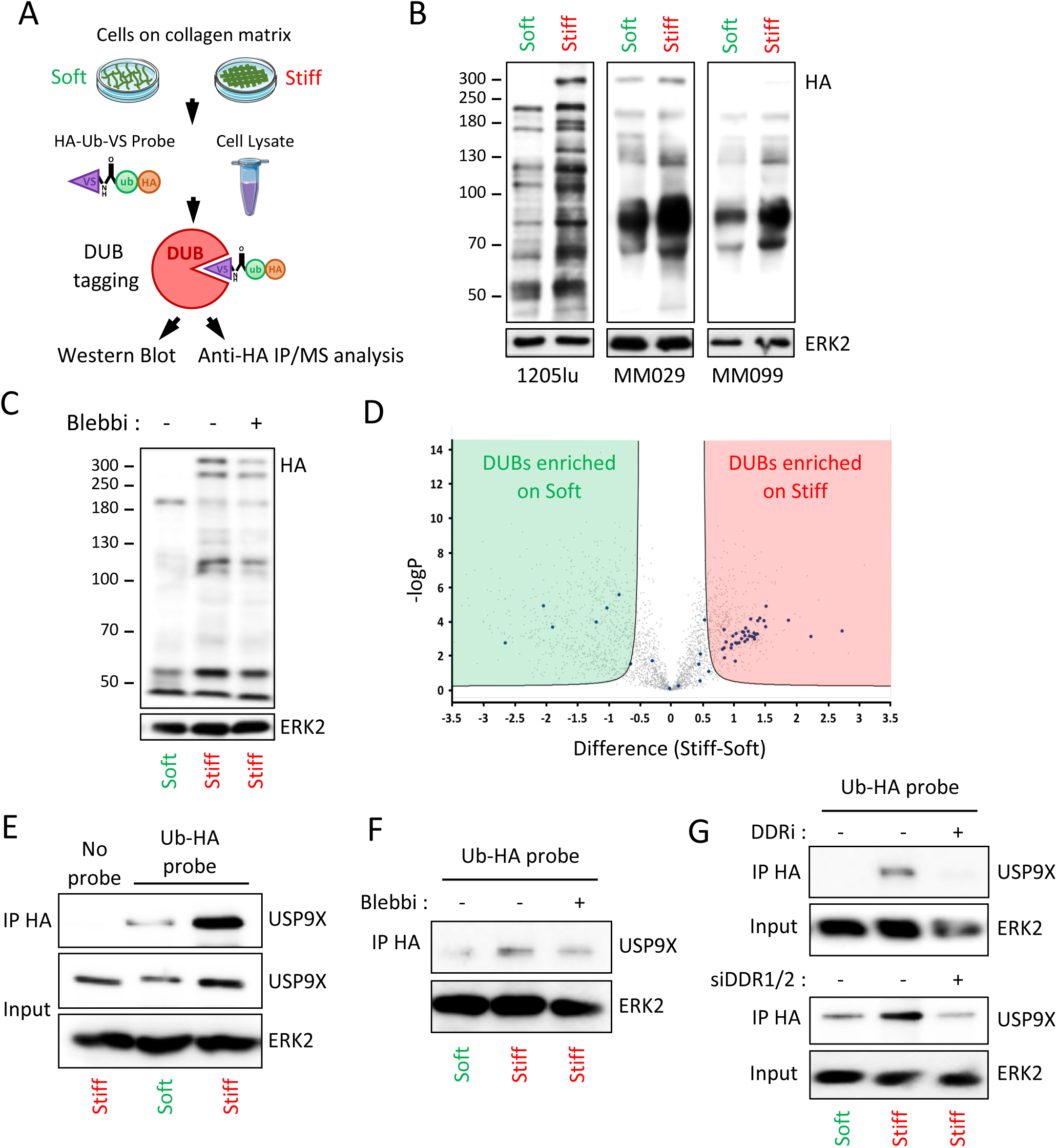
USP9X activity is controlled by extracellular collagen matrix stiffness. **A.** Principle of the in vitro labeling method of DUBs by the activity-based probe HA-Ub-VS (DUB trap assay) according to the collagen matrix stiffness. **B.** Effect of collagen matrix stiffness on DUBs activity in 1205Lu, MM029 and MM099 cells cultivated for 72h on soft versus stiff collagen matrices. Lysates of the indicated cells were incubated at 37°C with the HA-Ub-VS probe and analyzed by anti-HA. ERK2, loading control. **C.** Effect of myosin II inhibition using 20µM of Blebbistatin (Blebbi) on DUB activity according to collagen matrix stiffness on 1205Lu cells cultivated for 72h on soft versus stiff collagen matrices. Lysates were incubated with the HA-Ub-VS probe and analyzed by anti-HA. ERK2, loading control. **D.** 1205Lu cells were cultivated and lysed according to the DUB trap assay protocol in (A). Lysate containing HA-Ub-DUB complexes were immunoprecipitated using anti-HA coated agarose beads and analysed by quantitative proteomics to determine DUBs whose activity is affected by stiffness. Volcano-plot shows DUBs (blue dots) enriched on soft (green side) or stiff (red side) collagen matrices. **E.** Effect of collagen stiffness on USP9X activity. 1205Lu cells were cultivated for 72h on soft versus stiff collagen matrices. Cell lysates labelled with or without the HA-Ub-VS probe were immunoprecipitated using anti-HA coated agarose beads and analyzed by anti-USP9X immunoblot. **F.** Effect of myosin II inhibition on USP9X activity according to collagen matrix stiffness. 1205Lu cells were cultivated for 72h on soft versus stiff collagen matrices in the presence or not of 20µM Blebbistatin (Blebbi). Lysates were incubated with the HA-Ub-VS probe, immunoprecipitated with anti-HA coated agarose beads and analyzed by anti-USP9X immunoblot. **G.** Targeting DDR1/2 prevents USP9X activation by stiff collagen. 1205Lu cells, treated with DDR inhibitor (Imatinib, 10µM) or transfected with siRNA targeting DDR1 and DDR2, were cultivated for 72h on soft or stiff collagen gels. Lysates were incubated with the HA-Ub-VS probe, immunoprecipitated with anti-HA coated agarose beads and analyzed by anti-USP9X immunoblot. ERK2, loading control.

### Physical and functional interaction between USP9X and YAP

The choice of USP9X as the primary subject of this study was motivated by several reasons. First, bioinformatic data predict that the oncoprotein and mechanotransducer YAP (19) is a partner of USP9X (Fig. 2A). Second, YAP protein levels were found stabilized by ECM stiffening on different melanoma cell lines (Fig. 2B). Third, *USP9X* mRNA expression is significantly increased in melanoma compared to benign skin lesions (Fig. 2C). Fourth, *in silico* analyses revealed a link between *USP9X* levels and the transcriptional signature associated with *YAP* in melanoma (Fig. 2D) and both *USP9X* and *YAP1* gene expression were associated with poor prognosis when expressed at high levels in melanoma patients (Fig. 2E). These observations support the hypothesis that USP9X is a mechano-sensitive DUB, which controls YAP expression and thus the functional program driven by ECM stiffness in melanoma cells.

**Figure 2.**
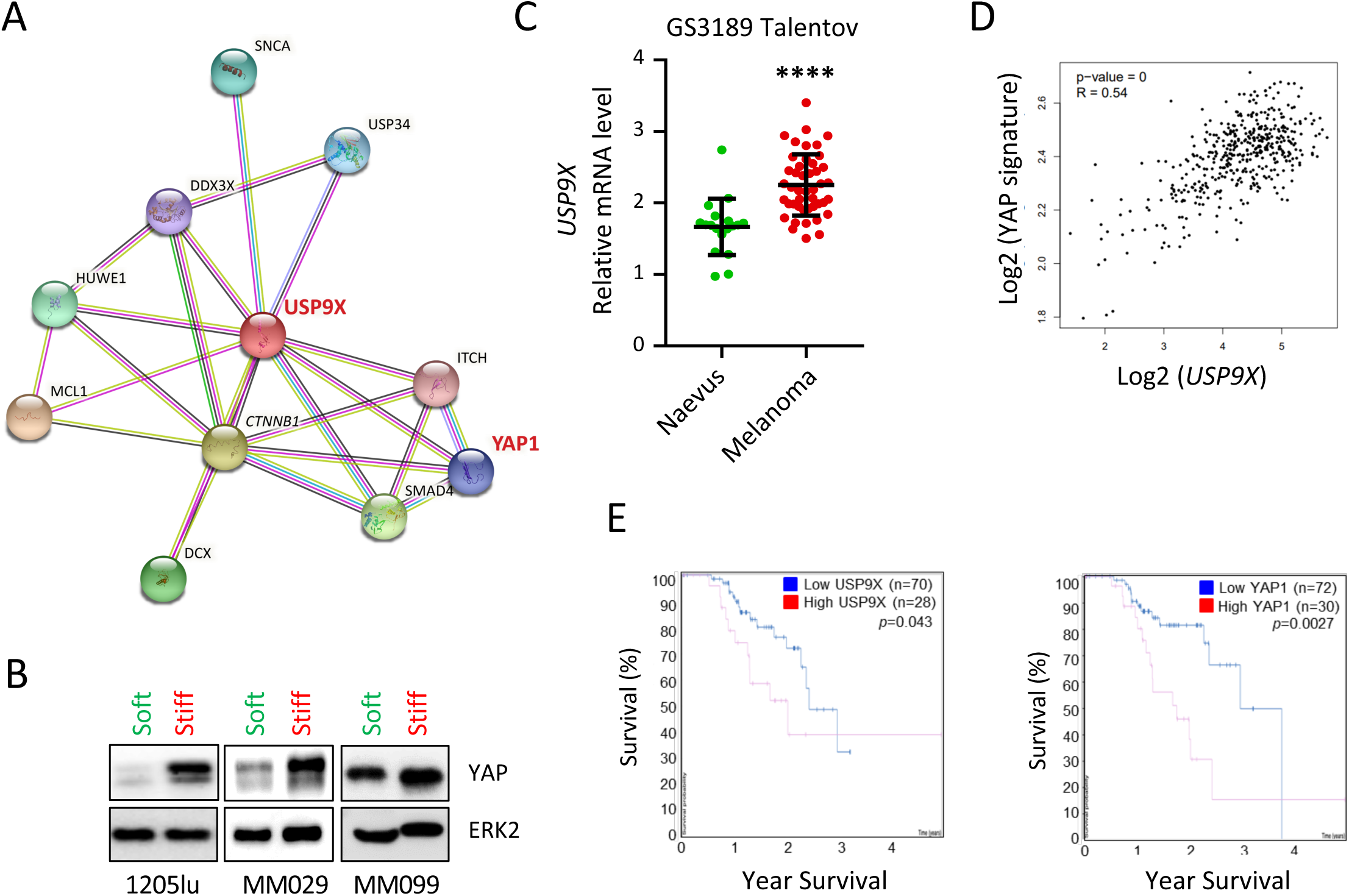
Correlation between USP9X and the YAP mechanosensor in melanoma. **A.** USP9X interactome generated using the STRING tool (https://string-db.org/network/9606.ENSP00000316357). **B.** Western blot analysis of YAP expression on 1205Lu, MM029 and MM099 cells cultivated for 72h on soft versus stiff collagen matrices. ERK2, loading control. **C.** Scatter plots showing the expression of *USP9X* mRNA in benign skin lesions (naevus) vs. metastatic melanoma (GSE3189 Talentov). ****p<0.0001. **D.** Pearson’s correlation analysis of *USP9X* levels and the transcriptional signature controlled by YAP (CORDENONSI YAP conserved signature) from data of skin melanoma extracted from the TCGA. Pearson correlation (R) and p-value are indicated. **E.** Kaplan-Meier curves showing melanoma patient survival percentage through time according to *USP9X* (left) and *YAP* expression (right). Data were retrieved from the skin melanoma TCGA database using ProteinAtlas (proteinatlas.org). p-values are indicated (logrank test).

### USP9X controls YAP protein levels and activity

To test this hypothesis, we used siRNA sequences directed against USP9X transfected into melanoma cell lines. This approach showed that USP9X silencing induces a loss of YAP protein expression without affecting its messenger (Fig. 3A, B). In contrast, USP9X knockdown induced the loss of the messenger of several YAP target genes (Fig. 3C). YAP is a transcriptional co-activator whose activity depends on its subcellular localization. Consistently, biochemical subcellular fragmentation indicated that stiff collagen increases the nuclear localization of USP9X and YAP, compared to soft collagen in 1205Lu melanoma cells (Supplementary Fig. 2). In addition, knockdown of USP9X decreased the nuclear shuttling of YAP on stiff ECM (Fig. 3D). Similar to the findings from USP9X silencing, treatment of 1205Lu cells with the USP9X inhibitor G9 induced a decrease in YAP protein expression (Fig. 3E), without affecting mRNA levels (Fig. 3F). However, we observed a dose-dependent loss of expression of several YAP target genes (Fig. 3G). Finally, pharmacological inhibition of USP9X also induced a dose-dependent decrease in the nuclear localization of YAP (Fig. 3H). In addition, depletion of USP9X induced the loss of YAP expression in other human (A375) and murine (YUMM1.7) melanoma cell lines, as well as in two short-term melanoma cell cultures (MM029 and MM099) and in the BRAF inhibitor-resistant melanoma cell line M238R (Fig. 3I). Interestingly, we also found that the loss of USP9X was associated with decreased YAP protein levels in other cancer cell lines, including lung (A549), breast (MDA-MB-231) and pancreatic (PANC) cancer cell lines (Supplementary Fig. 3A). Together, these results indicate that USP9X controls expression of the mechanosensor YAP at protein level in cancer cells including melanoma cells.

**Figure 3.**
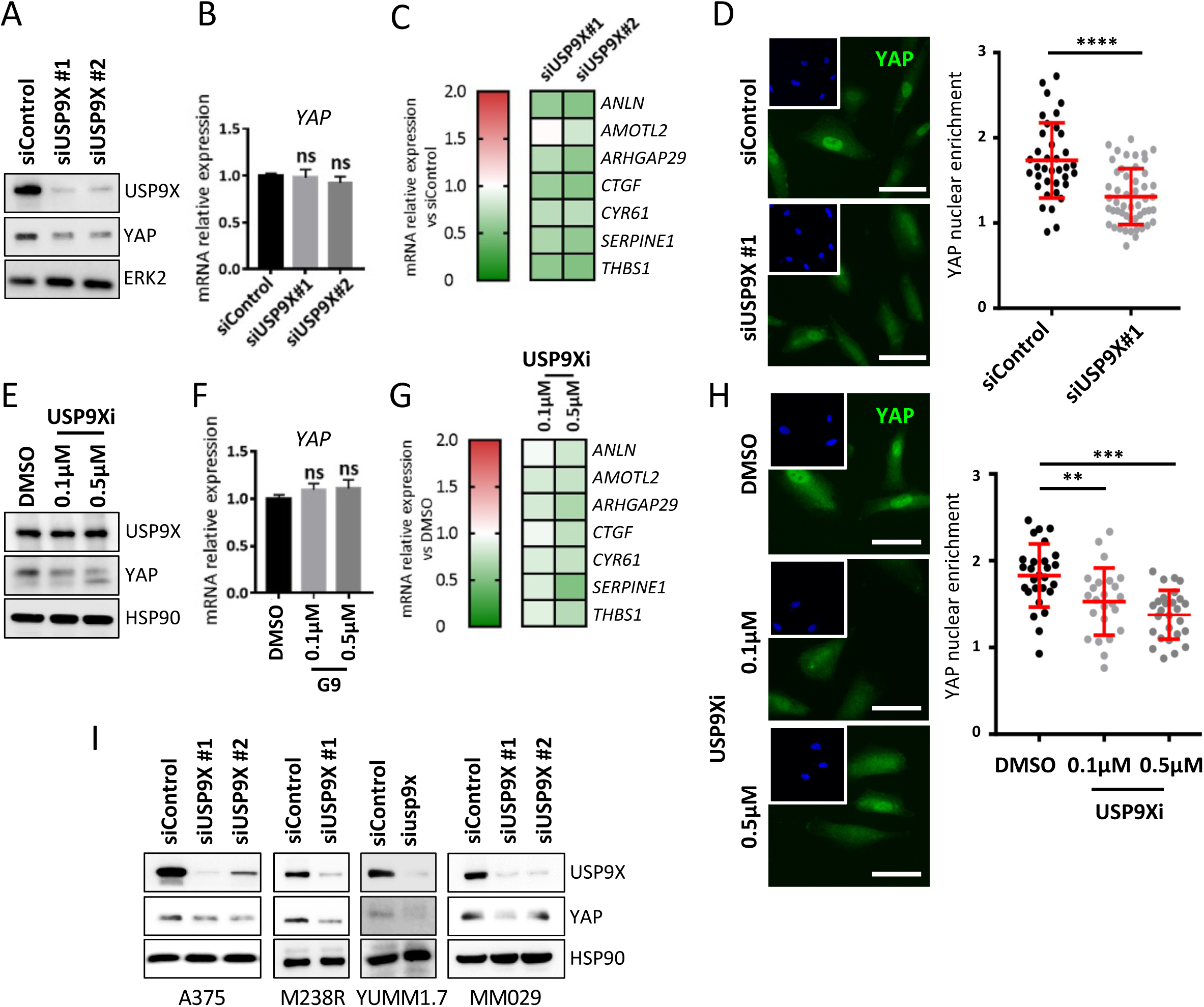
USP9X controls YAP expression and activity in melanoma. **A.** Western blot analysis of USP9X and YAP expression on 1205Lu cells transfected for 72h with 2 different sequences of siUSP9X or a control siRNA and cultivated on stiff collagen matrices. ERK2, loading control. **B.** Bar graphs showing the expression of *YAP* mRNA on cells transfected and cultivated according to (A). Data represent mean ± SD. ns, non-significant. **C.** Heatmap showing the expression of YAP target genes on 1205Lu cells transfected with 2 siUSP9X sequences or a control siRNA and cultivated on stiff collagen matrices as in (A). Data represents mRNA expression relative to the siControl condition. **D.** *Left,* Effect of USP9X depletion on YAP (green) nuclear translocation assessed by immunofluorescence in cells cultured for 72h on stiff collagen substrates. Insets show nuclei in blue. Scale bar, 50µm. *Right*, scatter plots showing the quantification of YAP nuclear enrichment in siControl and siUSP9X transfected cells. Data represent mean ± SD (n ≥ 30 cells per condition). Data are representative of 3 independent experiments. ****p<0.0001, Kruskal-Wallis analysis. **E.** Western blot analysis of USP9X and YAP expression on 1205Lu cells treated with 2 doses of USP9X inhibitor G9 for 72h and cultivated on stiff collagen matrices. ERK2, loading control. **F.** Bar graphs showing the expression of *YAP* mRNA on cells treated and cultivated according to (E). Data represent mean ± SD. ns, non-significant. **G.** Heatmap showing the expression of YAP target genes on 1205Lu cells treated with 2 doses of USP9X inhibitor G9 for 72h and cultivated on stiff collagen matrices as in (E). Data represents mRNA expression relative to the siControl condition. **H.** *Left,* immunofluorescence analysis of YAP (green) nuclear translocation in cells treated with USP9X inhibitor (G9) as in (E) and cultured for 72h on stiff collagen. Insets show nuclei in blue. Scale bar, 50µm. *Right*, scatter plots showing the quantification of YAP nuclear enrichment in control and G9-siControl and USP9Xi-treated cells. Data represent mean ± SD (n ≥ 30 cells per condition). Data are representative of 3 independent experiments. ***p<0.001, **p<0.01, Kruskal-Wallis analysis. **I.** Western blot analysis of USP9X and YAP expression on A375, YUMM1.7, M239R and MM029 cells transfected for 72h with different sequences of siUSP9X or a control siRNA and cultivated on stiff collagen matrices. ERK2, loading control.

### USP9X regulates the ubiquitination and degradation of YAP

We next examined how USP9X controls YAP protein levels. When USP9X was overexpressed in HEK293FT cells, an increase in YAP protein expression was observed that was not found with the overexpression of the catalytically inactive USP9X mutant (USP9X MT C1566A) (Fig. 4A). Previous studies showed that the E3 ligase β-TRCP ubiquitinates YAP after phosphorylation of its phosphodegron motif, leading to degradation by the proteasome, as part of the Hippo pathway (47). Here, we observed that depletion of β-TRCP in 1205Lu cells increased YAP protein expression (Fig. 4B) and nuclear subcellular location (Fig. 4C). In contrast, TAZ protein levels were not affected by β-TRCP silencing (Fig. 4B). To confirm that YAP is degraded at the proteasome level, we treated 1205Lu cells with the proteasome inhibitor Bortezomib (BTZ) for increasing times. As a control, the levels of the labile EMT factor SLUG (48) were also monitored. An increase in YAP expression after 4h of BTZ treatment was observed, indicating that YAP is degraded by the proteasome in melanoma cells (Fig. 4D). When these cells were treated with the protein synthesis inhibitor cycloheximide (CHX), a loss of SLUG expression but not YAP was found, suggesting that in contrast with SLUG, YAP is a relatively stable protein (Fig. 4E). Treatment with the USP9X inhibitor G9 had no effect on SLUG expression and only partially decreases YAP expression after 6h of treatment (Fig. 4F). However, the combination of USP9X inhibitor with CHX caused a total loss of YAP expression as early as 1h (Fig. 4G), suggesting that the deubiquitinase activity of USP9X protects YAP from degradation by preventing its addressing to the proteasome. To assess the regulation of YAP ubiquitination by USP9X, we used GST-TUBEs fusion proteins to pull-down poly-ubiquitinated proteins from treated melanoma cells (Fig. 4H). To prevent the degradation of poly-ubiquitinated proteins, cells were treated with BTZ. To prevent their possible deubiquitination by USP9X, cells were also treated or not with G9. Cell lysates were precipitated with GST-TUBEs and glutathione-agarose beads and analyzed by western blot. These experiments show that USP9X inhibition in the presence of BTZ increased YAP poly-ubiquitination, indicating that the deubiquitinase activity of USP9X is required to prevent YAP ubiquitination and degradation by the proteasome (Fig. 4I). In addition, we found that the expression of a proteasome-resistant form of MYC-tagged YAP, YAP 5SA was not affected by USP9X knockdown in melanoma cells in contrast to the endogenous form of YAP (Fig. 4J). This version of YAP is mutated on the 5 serine residues phosphorylated by LATS, which prevents its degradation and thus makes it constitutively active (49). Together, these results demonstrate that USP9X controls the ubiquitination and degradation of YAP in melanoma cells.

**Figure 4.**
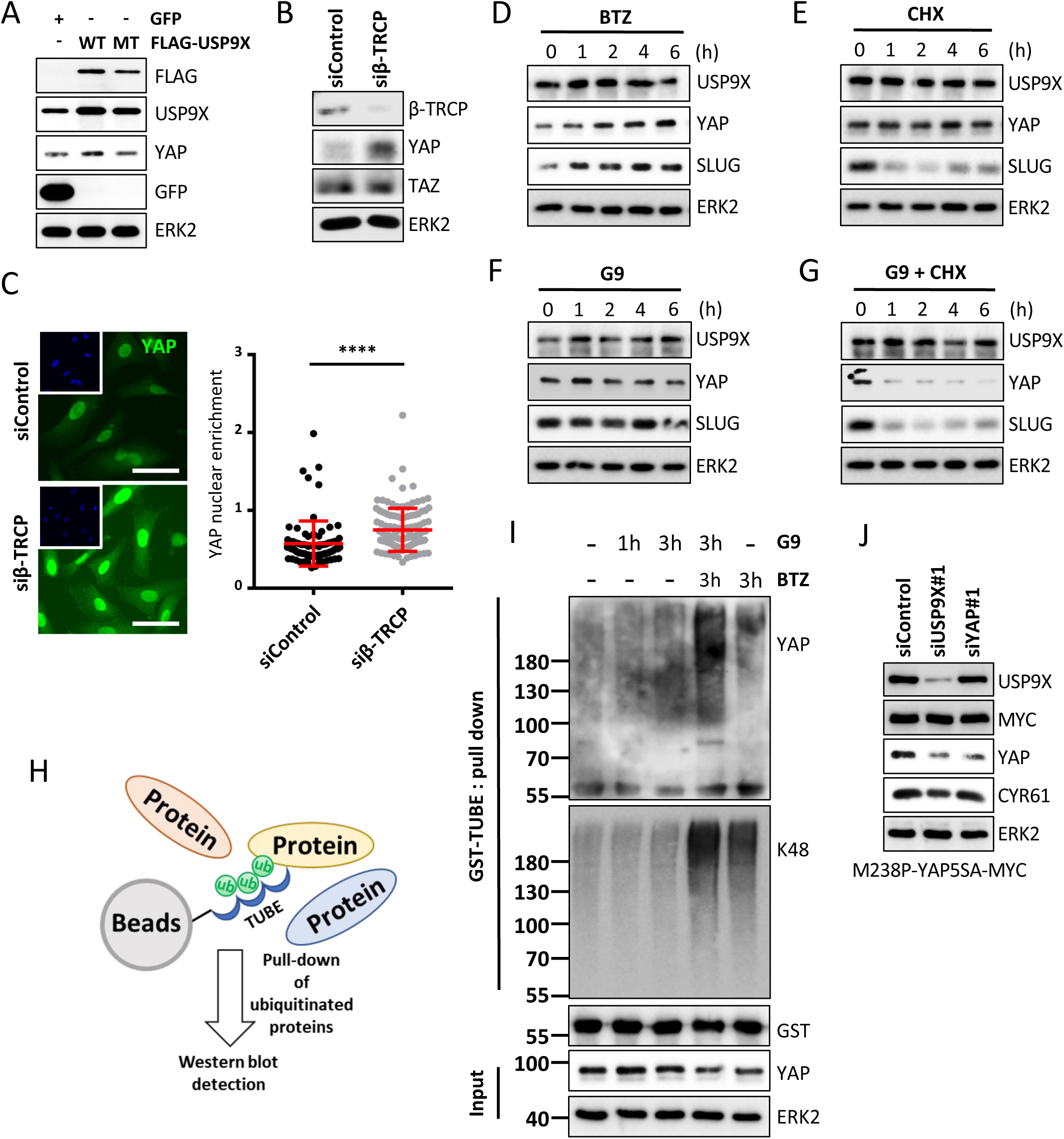
USP9X regulates YAP stability and prevents its proteasomal degradation. **A.** Western blot analysis of USP9X, FLAG, YAP and GFP expression on HEK293FT cells transfected for 48h with plasmid coding for a WT FLAG-USP9X, a dead catalytic mutant FLAG-USP9X or GFP as control. ERK2, loading control. **B.** Western blot analysis of YAP, TAZ and β-TRCP expression on 1205Lu cells transfected with a siβ-TRCP and a control siRNA for 72h and cultivated on stiff collagen matrices. ERK2, loading control. **C.** *Left,* Effect of β-TRCP depletion on YAP (green) nuclear translocation assessed by immunofluorescence in cells grown for 72h on stiff collagen matrices. Insets show nuclei in blue. Scale bar, 50µm. *Right*, scatter plots showing the quantification of YAP nuclear enrichment in siControl and siβ-TRCP transfected cells. Data represent mean ± SD (n ≥ 30 cells per condition). Data are representative of 3 independent experiments. ****p<0.0001, Kruskal-Wallis analysis. **D-G.** Western blot analysis of USP9X, YAP and SLUG on 1205Lu cells treated for the indicated times with 100nM of the proteasome inhibitor Bortezomib (BTZ) **(D),** 25µg/ml of the protein synthesis inhibitor Cycloheximide (CHX) **(E),** 1µM of the USP9X inhibitor G9 **(F),** or with a combination of Cycloheximide (25µg/ml) and G9 (1µM) **(G)**. ERK2, loading control. **H.** Principle of the affinity-based precipitation of poly-ubiquitinated proteins using GST-TUBE (Tandem Ubiquitin Binding Entities) fusion protein and glutathione coated agarose beads. **I.** GST-TUBE pull down analysis of YAP ubiquitination on 1205Lu cells treated for 3h with G9 (1µM), BTZ (100nM) or a combination of both. Precipitated proteins were visualized using anti-K48 ubiquitin chain antibody and anti-YAP antibody. ERK2, loading control. **J.** Western Blot analysis of USP9X, YAP, MYC and CYR61 expression on M238P melanoma cells overexpressing MYC tagged YAP5SA constitutive active mutant transfected with siUSP9X#1, siYAP#1 or a control siRNA and cultivated for 72h on stiff collagen matrices. ERK2, loading control.

### USP9X promotes melanoma cell migration and invasion *in vitro* and *in vivo*

Besides its role as mechanotransducer, YAP is involved in the migratory and invasive capabilities of cutaneous melanoma cells (50). To examine the functional link between USP9X and YAP in melanoma, we performed a scratch assay in the invasive melanoma cells 1205Lu transfected with two different interfering RNA sequences directed against USP9X. Depletion of USP9X impaired cell migration on stiff substrate (Fig. 5A), as was observed after depletion of YAP (Fig. 5B). Consistently, random motility (Fig. 5C) and chemotactic migration (Fig. 5D) of USP9X-depleted cells cultured on stiff collagen substrate was significantly decreased compared to control cells. These data were confirmed by pharmacological treatment with the USP9X inhibitor, which significantly and dose-dependently decreased motility and chemotactic migration of 1205Lu cells (Fig. 5E, F). Interestingly, depletion of USP9X also induced a decrease in motility in a panel of solid cancer cell lines (Supplementary Fig. 3B). Finally, overexpression of a proteasome-resistant YAP mutant (YAP5SA) in M238P melanoma cells rescued cell motility that was inhibited by USP9X and YAP depletion (Fig. 5G).

**Figure 5.**
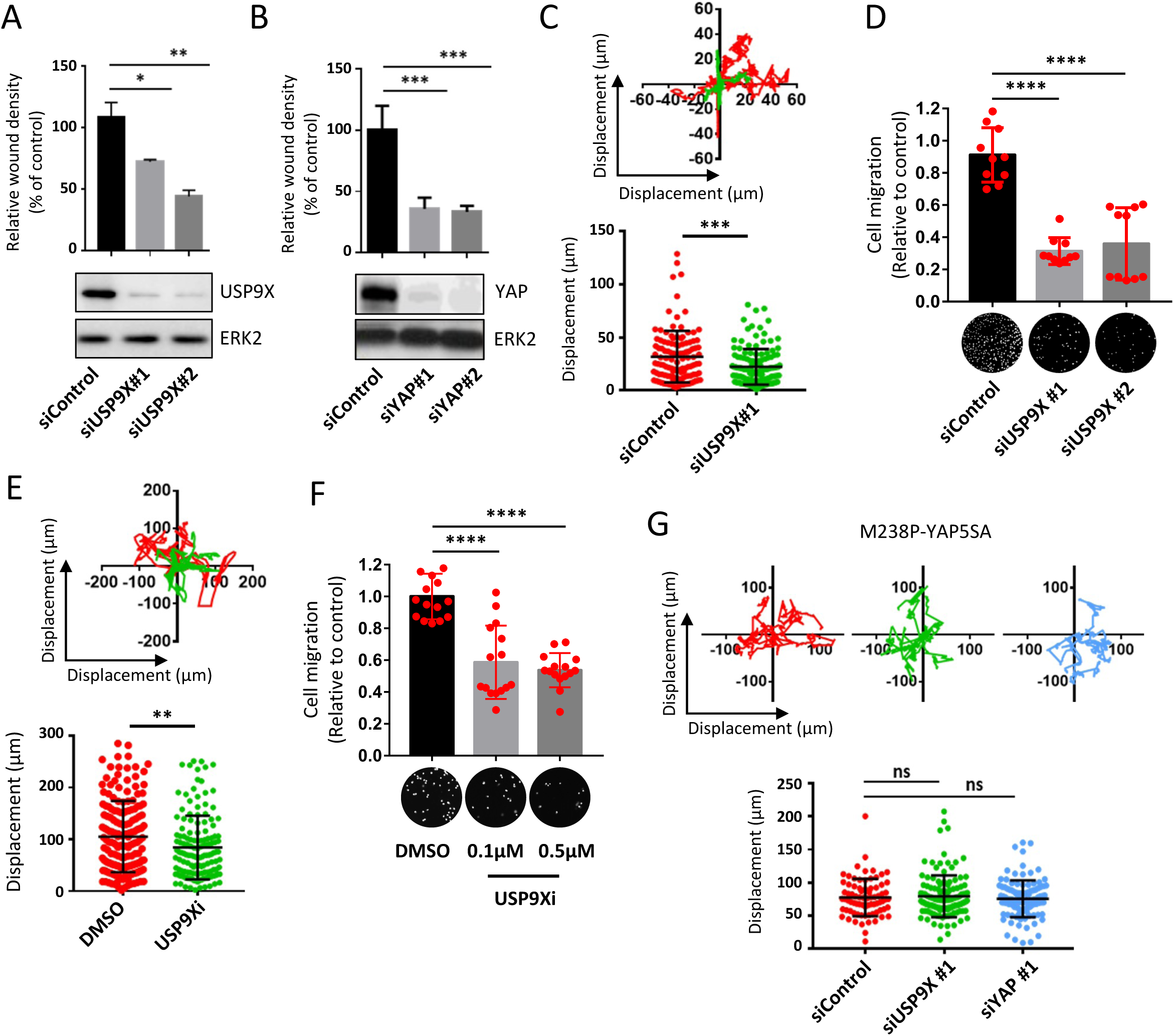
USP9X promotes melanoma cell migration through YAP stabilization. **A-B.** Motility of 1205 cells on stiff collagen matrices was assayed by scratch assays following transfection with control siRNA, 2 different sequences of siUSP9X **(A)** or 2 different sequences of siYAP **(B)**. Confluent transfected cell monolayers were wounded and imaged after 24 h. Migrating cells in scratch area were quantified relative to the siControl condition (*upper panels*). *Bottom panels*, western blot analysis of USP9X and YAP on transfected cells. ERK2, loading control. Bar graphs represent means ± SD (n = 3 independent experiments). ***p< 0.001, **p< 0.01, *p< 0.05, two-way ANOVA and Bonferroni post-tests. **C.** Time-lapse microscopy analysis of 1205Lu cells displacement on stiff collagen after siUSP9X#1 or siControl transfection. Cell migration was recorded over 24h and automatically tracked using Image J. *Upper panel*, displacement path of 5 representative cells for each condition. *Lower panel*, scatter plot showing the quantification of cell displacement (n>40). Data represent median ± quartiles. *** p<0.001, Kruskal-Wallis analysis. **D.** Bar graphs showing the analysis of chemotactic migration in Boyden chambers of 1205Lu cells transfected with siUSP9X sequences or a control siRNA. Representative images (*bottom*) and quantification (*top*) of migrating cells in the different conditions. Data represents the mean ± SD of 3 independent experiments. ****p<0.0001, Kruskal-Wallis analysis. **E.** Time-lapse microscopy analysis of 1205Lu cells displacement on stiff collagen after after treatment with the USP9X inhibitor G9 (1µM). *Upper panel*, displacement path of 5 representative cells for each condition. *Lower panel*, scatter plot showing the quantification of cell displacement (n>40). Data represent median ± quartiles. ** p<0.01, Kruskal-Wallis analysis. **F.** Quantification of chemotactic migration of 1205Lu cells treated with increasing doses of G9. Representative images (*bottom*) and quantification (*top*) of migrating cells in the different conditions. Data represents the mean ± SD of 3 independent experiments. ****p<0.0001, Kruskal-Wallis analysis. **G.** Time-lapse microscopy analysis of M238P-YAP5SA cell displacement during 24h after siUSP9X#1, siYAP#1 or siControl transfection. *Upper panel*, displacement path of 5 representative cells for each condition. *Lower panel*, scatter plot showing the quantification of cell displacement (n>40). Data represent median ± quartiles. ns, non-significant, Kruskal-Wallis analysis.

To study the implication of USP9X in melanoma cell invasion, we generated lentiviral vectors containing a pSGEP Luc-PuroR plasmid expressing different shUSP9X sequences or a non-targeting shRNA sequence (directed against the Renilla luciferase mRNA). We generated 1205Lu cells stably depleted for USP9X using these lentiviral vectors and analyzed their invasive properties in stiff collagen conditions. First, we confirmed that stable depletion of USP9X prevented the upregulation of YAP induced by collagen stiffening (Fig. 6A). Consistently, we noticed a loss of YAP nuclear localization in USP9X stably depleted cells cultured on stiff collagen matrix (Fig. 6B). Interestingly, we observed that YAP protein expression was strongly reduced in USP9X-depleted cells cultured on soft collagen compared to control cells (Fig.6A). Cells stably depleted for USP9X also showed a significant decrease in motility when plated on a stiff collagen substrate (Fig. 6C). Notably, the motility of USP9X depleted cells seeded on stiff collagen was not statistically different from that of shControl cells seeded on a soft collagen (Fig. 6C), suggesting that the regulation of YAP by USP9X is important for mechanosensing. In addition, stable depletion of USP9X significantly decreased melanoma cell invasiveness when grown as spheroids in a 3D collagen/matrigel matrix (Fig. 6D).

**Figure 6.**
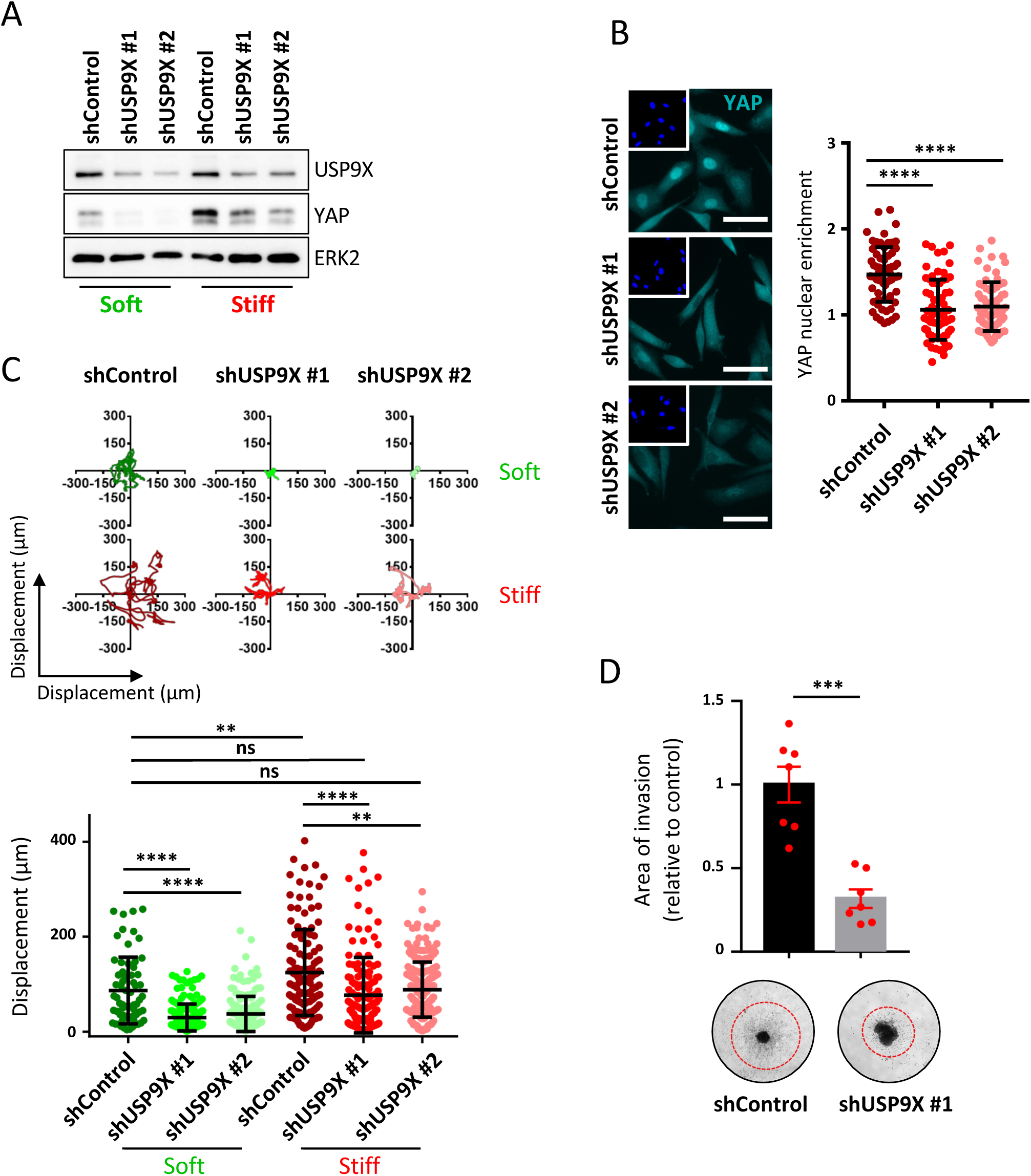
Stable depletion of USP9X impairs collagen stiffness-induced melanoma cell migration and invasion. **A.** Western blot analysis of USP9X and YAP expression on 1205Lu cells stably depleted of USP9X and cultivated for 72h on soft versus stiff collagen matrices. ERK2, loading control. **B.** *Left*, Representative Images of YAP staining (Cyan) on 1205Lu cells stably depleted of USP9X and seeded on stiff collagen substrates. Insets show nuclei stained with Hoechst (Blue). Scale bar, 50µm. *Right*, Quantification of the nucleocytoplasmic distribution of YAP (n ≥ 30 cells per condition). Data represent mean ± SD. Data are representative of 3 independent experiments. ****, P<0.0001, Kruskal-Wallis analysis. **C.** Time-lapse microscopy analysis of shControl, shUSP9X #1 and shUSP9X #2 cell displacement during 24h on soft vs. stiff collagen matrices. *Upper panel*, displacement path of 5 representative cells for each condition. *Lower panel*, quantification of cell displacement (n>40). ****p<0.0001, **p<0.01, ns, non-significant, Kruskal-Wallis analysis. **D.** Analysis of cell invasion of shControl and shUSP9X #1 cells grown in spheroids and included in 3D collagen matrices. *Lower panel*, representative images of cell invasion after 24h (dotted red circles). *Upper panel,* quantification of cell invasion area relative to control. *** P<0.001, Kruskal-Wallis analysis.

We next injected shControl and shUSP9X#1 bioluminescent cells into the tail vein of nude mice to monitor melanoma cell extravasation into lungs and long-term metastatic burden. Bioluminescence imaging recorded after 3h of tail vein injection revealed a decreased ability of USP9X-depleted cells to extravasate into the lungs (Fig. 7A). A second series of *in vivo* bioluminescence monitoring was performed every 7 days until the end of the experiment. (56 days). When cells were stably depleted for USP9X, no lung metastases were observed *in vivo* and *ex vivo* (Fig. 7B, C). Together, these data suggest that USP9X targeting affects YAP-dependent tumor cell migration and invasion on stiff conditions.

**Figure 7.**
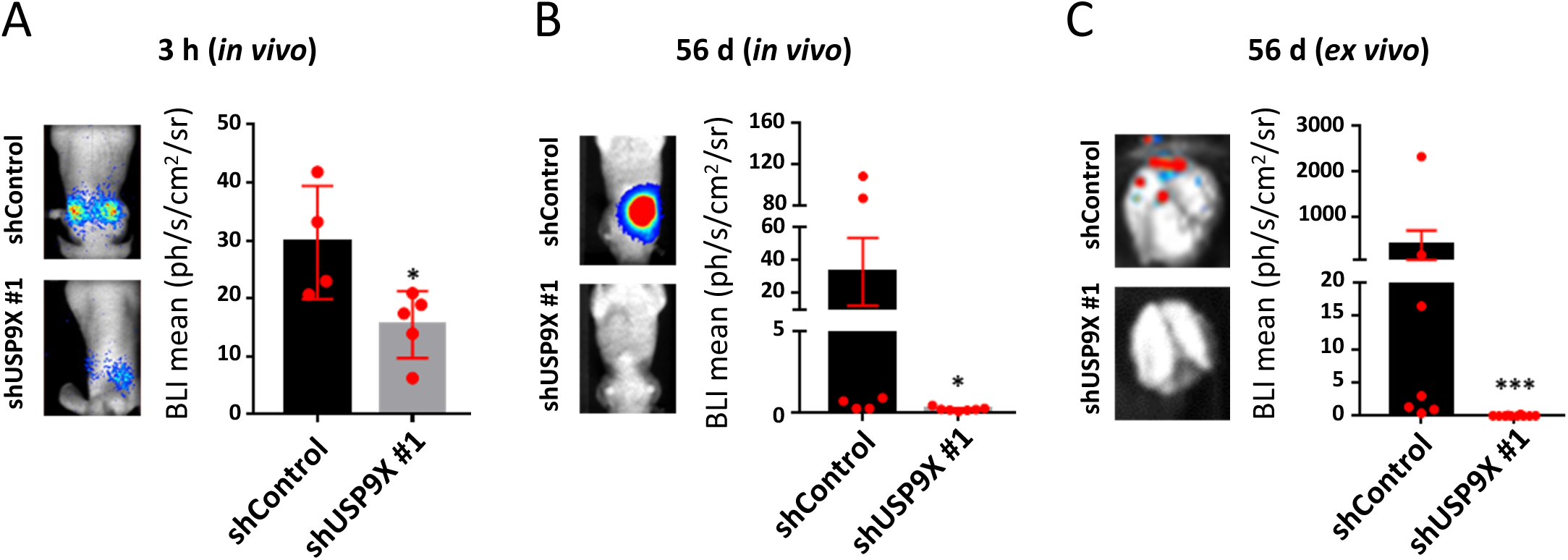
USP9X promotes melanoma invasiveness *in vivo.* **A.** shControl and shUSP9X #1 cells were injected by tail vein into nude mice and cell extravasation into lungs was monitored using a photon imager 3h after injection. *Left*, representative images of photon fluxes produced by bioluminescent melanoma cells. *Right*, normalized photon flux (BLI) of lungs from control (shControl, n = 4) and USP9X-depleted (shUSP9X, n = 5) injected mice. *p<0.05, unpaired two-tailed Mann–Whitney’s test. **B.** shControl and shUSP9X #1 cells were injected by tail vein into nude mice and lung metastasic progression was monitored and quantified using a photon imager. *Left*, representative images of photon fluxes produced by bioluminescent melanoma cells at the end of the experiment (56 days). *Right*, normalized photon flux (BLI) of lungs from control (shControl, n = 6) and USP9X-depleted (shUSP9X, n = 7) injected mice. *p<0.05, unpaired two-tailed Mann–Whitney’s test. **C.** Quantification of lung metastatic foci by ex vivo BLI at the endpoint of the extravasation assay performed in (B). Representative images of lung metastases (left) and quantification of lung metastatic foci per lung (righ). ***p<0.001, unpaired two-tailed Mann–Whitney’s test.

### Targeting USP9X prevents collagen network remodeling induced by targeted therapies and delays treatment relapse

YAP plays a central role in ECM remodeling, mechanical adaptation and resistance of melanoma cells to targeted therapy (22,35,41). We first assessed the contribution of USP9X in melanoma cell contractility induced by mutant BRAF inhibition. Collagen contraction assays of 1205Lu cells pretreated with the BRAF inhibitor (BRAFi) in the presence or not of G9 revealed that pharmacological inhibition of USP9X impaired cell contractility induced by oncogenic BRAF pathway inhibition (Supplementary Fig. 4A). Time-lapse cell imaging next showed that treating 1205Lu cells with G9 in combination with BRAF/MEK inhibitors (BRAFi/MEKi) dramatically decreased cell proliferation and induced caspase 3 cleavage compared to what was observed with BRAFi/MEKi alone (Supplementary Fig. 4B, C). Of note, G9 as a single agent had no effect compared to control DMSO on cell proliferation monitored up to 5 days. These data suggest that YAP negative regulation downstream USP9X inhibition enhances the cytotoxic effect of the targeted therapy.

We next investigated whether inhibition of USP9X affects the fibrotic-like reaction induced by BRAF-targeted therapy and drug resistance. To test this, the activity of G9 combined with BRAFi/MEKi was assessed in a pre-clinical syngeneic melanoma model. Murine BRAF-mutated melanoma cells YUMM1.7 were injected subcutaneously into C57BL/6 mice, which were treated with G9, BRAFi/MEKi or the triple combination of BRAFi/MEKi plus G9 (Fig. 8A). G9 did not display significant anti-melanoma effect when administered alone, slightly slowing down tumor growth but not triggering tumor volume decrease (Fig. 8B). After BRAFi/MEKi administration, melanomas that had initially responded to the targeted therapy rapidly relapsed and resumed growth in most tumors. In contrast, combination of BRAFi/MEKi and G9 significantly delayed tumor relapse (Fig. 8B) and improved mouse survival (Fig. 8C). Immunohistochemical analysis of tumor samples further documented that in comparison to single regimen, the combined BRAFi/MEKi and G9 treatment dramatically reduced the dense ECM remodeling induced by targeted therapy as shown by polarized light imaging on Sirius red labeled tumor slides (Fig 8D), quantification of collagen fibers area (Fig 8E) and second harmonic generation (SHG) imaging of collagen fibrillar network (Fig. 8F). Consistently, transcriptomic analysis of tumor samples showed that the triple combination BRAFi/MEKi/G9 reduced the expression of YAP target genes Ctgf, Thbs1 and Myl9 that were triggered by the targeted treatment (Fig. 8G). These data indicate that USP9X inhibition counteracted the fibrotic-like reaction induced by targeted therapy in melanoma and delayed tumor relapse.

**Figure 8.**
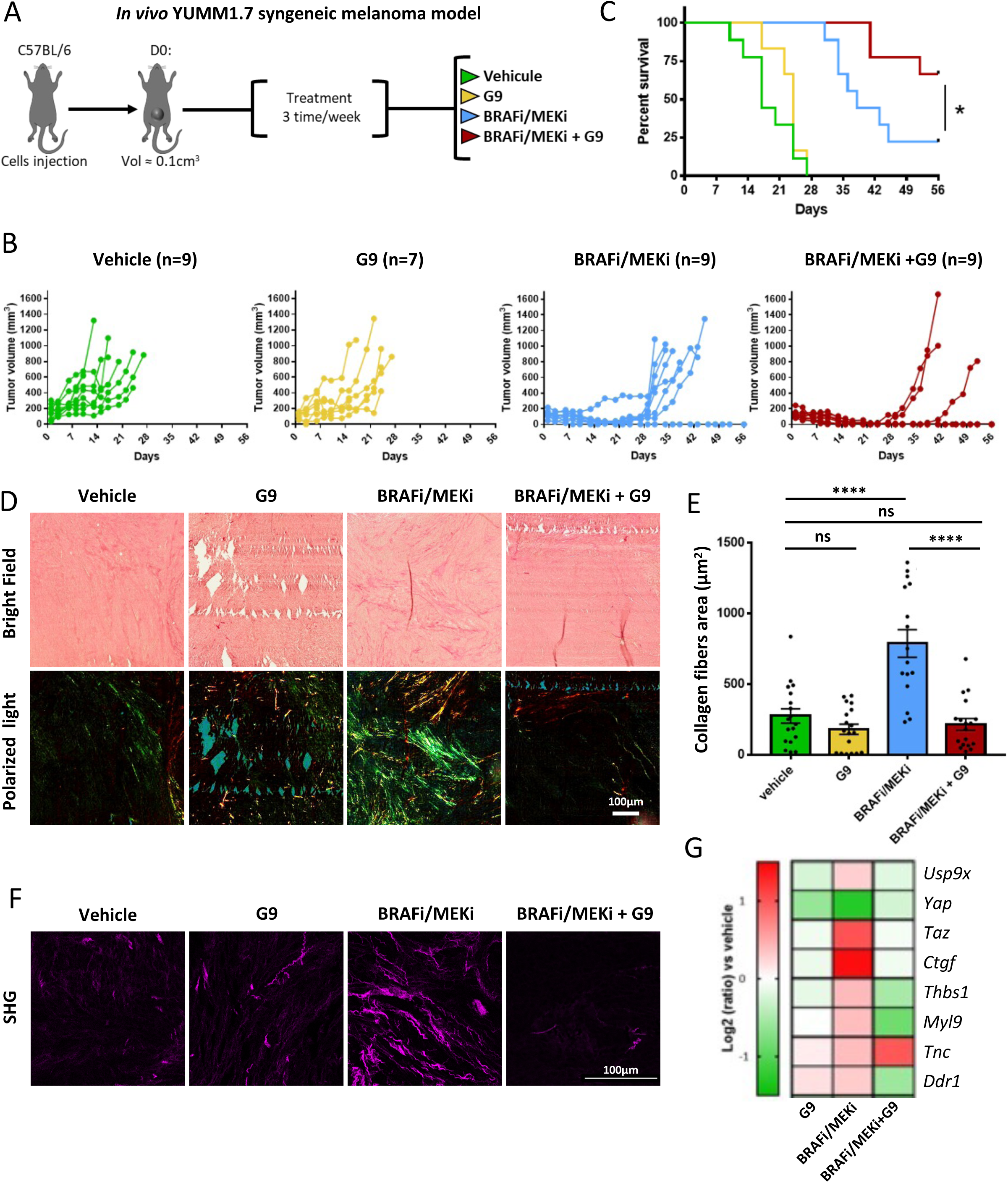
Targeting USP9X prevents collagen network remodeling induced by targeted therapies and delays treatment relapse. **A.** Schematic representation of the experimental procedure used in this study. YUMM1.7 murine melanoma cells were subcutaneously injected in C57BL/6 mice. Once tumors reached 100mm^3^, mice were treated with vehicle, USP9X inhibitor G9 (7.5 mg/kg i.p.), BRAFi/MEKi (vemurafenib 30 mg/kg p.o./Trametinib 0.3 mg/kg p.o.), or the combination of BRAFi/MEKi and G9. **B.** Curves show tumor growth across time on individual animal within each treatment groups. **C.** Kaplan–Meier survival curves of mice treated with the indicated therapies. The log rank (Mantel–Cox) statistical test was used for BRAFi/MEKi vs BRAFi/MEKi/G9. *p<0.05. **D.** Representative images of the network of collagen fibers in tumors from mice under the different treatments. Tumor sections from the experiment shown in (C) were stained with picrosirius red and imaged under bright field and polarized light microscopy. Scale bar, 100µm. **E.** Collagen fibers area was quantified from picrosirius red stainings with ImageJ. Bar graph shows the mean ± SD of > 12 independent fields. ****p<0.0001, ns, non-significant, Kruskal-Wallis analysis. **F.** Second harmonic generation (SHG) microscopy on samples from (D). Scale bar = 100µm. **G.** Heatmap showing the differential expression of USP9X, YAP, TAZ, YAP target genes and ECM genes in untreated versus G9-, BRAFi/MEKi- or BRAFi/MEKi+G9-treated tumors. Gene expression was assessed by RT-qPCR.

## Discussion

Tumors are characterized by abnormal ECM deposition and greater stiffness than healthy tissue, which correlates with aggressiveness (17). Mechanical stresses that stimulate tumor cell proliferation, survival and invasion, also promote angiogenesis, hypoxia, and decrease the anti-tumor immune response. ECM stiffening is also involved in therapeutic escape, and we recently showed that melanoma cell mechanical plasticity in response to MAPK pathway inhibition triggers a YAP-dependent pro-fibrotic loop and tumor stiffening, involved in drug adaptation and resistance (22,35,41). Together with the ubiquitin-proteasome system, DUBs represent essential gatekeepers of protein homeostasis and, as such, play essential roles in virtually all biological processes (8,15). However, whether DUBs contribute to ECM biomechanical responses is still unknown.

To address this question, we implemented a screening strategy based on melanoma cells cultivated on collagen matrices with various stiffnesses combined to an activity-based ubiquitin probe for profiling DUB activity (44) and quantitative proteomics. We identified DUBs whose activity is modulated by collagen matrix stiffness, including the ubiquitin-specific protease 9X (USP9X) (also known as FAM), a highly conserved DUB from Drosophila to mammals (51). USP9X has been described before in melanoma in the regulation of NRAS through its stabilizing activity of the transcription factor ETS-1 (45) and in drug resistance via its effect on SOX2 (52). USP9X is also an important player in the migration of neuronal cells (53), which share with melanocytes the same embryonic origin (54). USP9X is also a key regulator of TGF® signaling (19), a pro-tumorigenic pathway in cutaneous melanoma (55). Interestingly, no studies have linked mechanotransduction and USP9X in cancer.

*In silico* analysis showed that the mechanotransducer YAP belongs to the USP9X interactome. This observation is consistent with a study defining YAP as the substrate of USP9X in breast cancer (56). Furthermore, in the SKCM cohort from the TCGA, USP9X expression correlates with the transcriptional signature of YAP target genes and both proteins are found associated with a poor prognosis in melanoma. In line with the mechanical plasticity of dedifferentiated invasive and drug resistant melanoma cells (22), YAP protein expression is increased in response to stiff ECM in several invasive melanoma cells. The literature also described the importance of matrix stiffness in migration and invasion in breast cancer (16) and the involvement of YAP and its transcription factor TEAD in melanoma migration (50,57). We therefore hypothesized that mechanical activation of USP9X by stiff ECM stabilizes YAP and controls its action on melanoma invasion and resistance. Interestingly, the effect induced by extracellular mechanical stress on the global activity of DUBs, including USP9X, is found dependent on the actomyosin cytoskeleton and myosin II activity. The collagen receptors DDR are also identified as the mechanosensors of collagen stiffness that control USP9X activity in melanoma cells. Given the role of myosin II in DDR1-mediated collagen contraction (46), it is likely that the activation of USP9X (and possibly of other DUBs) by mechanical signals is controlled by a direct interaction between DDR and myosin II. In line with the findings of our laboratory that DDR1 and DDR2 contribute to ECM-mediated drug resistance in melanoma (36), it would be relevant to study how more physiological settings such as cell-derived 3D ECM affect DUB activity in cancer cells and in other mechanically competent stromal cells, including fibroblasts or macrophages. Whether other major ECM receptors, like Integrins (17,58), mechanically regulate DUB activity remains an open question.

We next show that depletion or pharmacological targeting of USP9X causes loss of YAP protein expression without affecting its messenger. This observation is consistent with a post-translational stabilization of YAP by USP9X via its deubiquitinase activity. This hypothesis is supported by the observation that overexpression of the wild-type version of USP9X induces an increase in the protein level of YAP that is not found with its catalytically inactive mutant. A loss of nuclear localization of YAP and of its activity was also observed, consistent with a reduction of the expression of YAP/TEAD target genes. The relationship between USP9X and YAP is also confirmed *in silico* in the TCGA melanoma patient cohort, in other human and murine melanoma cell lines as well as in short-term primary cultures of melanoma. Importantly, USP9X also regulates YAP protein expression in other cancers including breast, pancreas and lung carcinomas, where ECM-dependent mechanical signals are known to foster tumor growth and progression.

When the Hippo pathway is active, YAP is phosphorylated by LATS kinases which reveals its phosphodegron motif allowing its ubiquitination by β-TRCP and its degradation at the proteasome (47). Consistent with this mechanism, depletion of β-TRCP increases YAP expression and consequently its nuclear localization. This confirms that YAP can be ubiquitinated and sent to the proteasome for degradation in our model. Interestingly the YAP homolog TAZ is not impacted by β-TRCP depletion in melanoma cells. Pharmacological inhibition of the proteasome increases YAP expression, which again confirms its regulation by the UPS in melanoma cells. On the contrary the blockade of protein synthesis is not sufficient to induce a loss of short-term expression of YAP. However, the combination of protein synthesis blockade with USP9X inhibition causes a complete loss of YAP expression, revealing that USP9X activity protects YAP from degradation upon protein synthesis inhibition. An affinity purification approach for the polyubiquitinated protein fraction further confirmed that USP9X activity prevents the ubiquitination of YAP and thus its degradation by the proteasome. In addition, we show that USP9X depletion does not affect the expression and transcriptional activity of a proteasome-resistant YAP mutant. Taken together, our findings support the notion that USP9X activity is responsible for YAP stabilization. USP9X structure shows a nuclear localization sequence (51) and we observed that both USP9X and YAP translocate into the nucleus upon stiff collagen signals. A question that remains opened is whether YAP deubiquitination by USP9X occurs into the cytoplasm or the nucleus.

Subsequently, we show that targeting USP9X phenocopies YAP-mediated functional outcomes, including cell migration, invasion and resistance to targeted therapy *in vitro* and *in vivo*. To determine that the effect of USP9X on melanoma cell migration is indeed YAP-dependent, we depleted USP9X in YAP5SA melanoma cells, showing that the non-degradable version of YAP rescues cell motility that is impaired in the absence of USP9X. Interestingly, the motility of USP9X-silenced melanoma cells grown on a stiff collagen matrix was not statistically different from that of control cells grown on a soft collagen matrix, supporting the idea that the loss of USP9X prevents cells from correctly interpret ECM mechanical signals. In line with a role of USP9X in cell migration and invasion that we observe in this study, we provide evidence that USP9X also contributes to melanoma cell extravasation and lung metastasis.

Finally, we sought to understand the implication of USP9X within the YAP-dependent mechanoresistance loop that we described (22). We show that USP9X inhibition increases the sensitivity of melanoma cells to oncogenic BRAF inhibition *in vitro* and impaired BRAFi-induced cell contractility. To extend these observations *in vivo*, we used a syngeneic model of melanoma response to BRAF and MEK inhibition (41). In this model, targeting USP9X counteracts the fibrotic-like response induced by the targeted therapy, including collagen fibers remodeling and expression of myofibroblastic markers and YAP target genes. Remarkably, the combination of USP9X inhibitor with BRAFi/MEKi enhances treatment efficacy and delays tumor relapse. Thus, our work provides further insights on the role of YAP in the mechanical feed-forward loop that takes place during non-genetic resistance to targeted therapies in melanoma and the involvement of the USP9X DUB in this process (22). Recent studies have underlined mechanisms that may be involved in cross-resistance between targeted therapies and immune checkpoint blockade therapies, including a YAP1 enrichment signature (59,60) and the activation of ROCK/actomyosin pathway (61). In this context, it would be interesting to investigate how the mechanically activated couple USP9X/YAP participates to therapeutic cross-resistance. Finally, the functional relationship between USP9X and YAP was observed in other cancer cell lines, suggesting that USP9X is also a regulator a YAP-dependent mechanical pathways in a broad array of solid tumors, including lung, breast and pancreatic carcinomas.

In conclusion, our work uncovers a novel regulation of the activity of the DUB USP9X by the mechanical microenvironment, resulting in stabilization of the key mechanotransducer YAP. This mechanosignaling is associated with increased melanoma invasive behavior and drug resistance. We therefore propose that USP9X is a “mechano-DUB” whose targeting could prevent the development of ECM-mediated progression and resistance of metastatic melanoma and provide novel therapeutic opportunities (Fig. 9).

**Figure 9.**
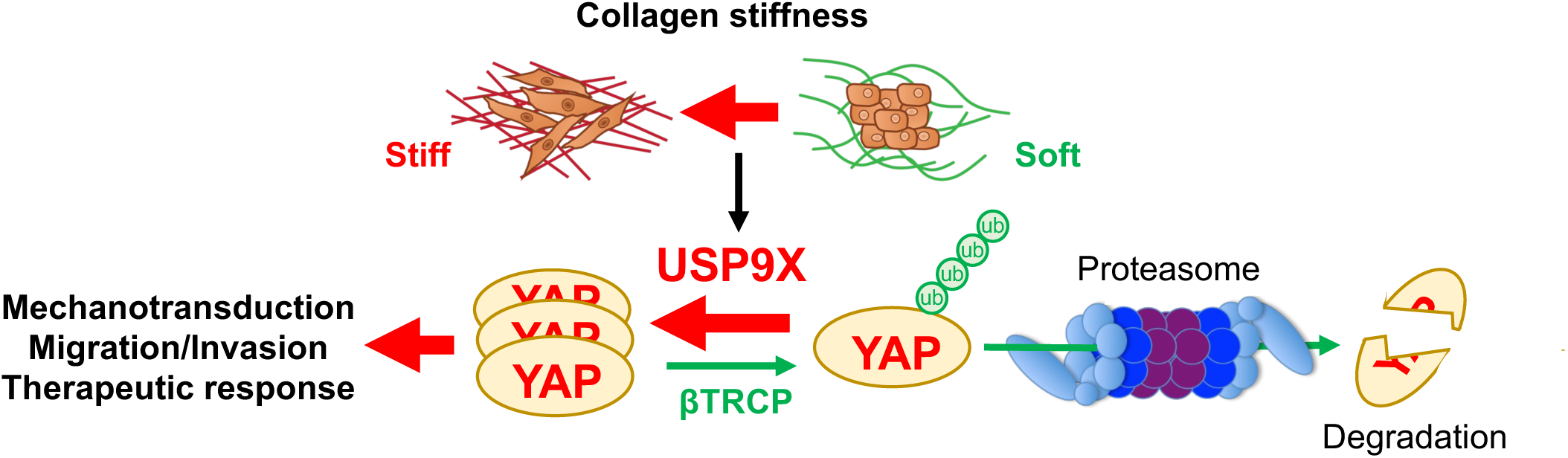
Schematic diagram of USP9X action in melanoma mechanobiology showing the relationship between collagen stiffness, USP9X and YAP-dependent mechanosignaling contributing to melanoma invasiveness and therapeutic resistance.

## Materials and Methods

### Cell culture and reagents

BRAF mutant melanoma cell lines 1205Lu and A375 were obtained as previously described (44,62). The isogenic pair of Vemurafenib-sensitive (P) and resistant (R) cell line M238 was described before (35,63). Melanoma cells were cultured in DMEM supplemented with 7% fetal bovine serum (FBS) (HyClone) and 1% penicillin/streptomycin (P/S) solution. Short-term cultures of patient melanoma cells MM099 and MM029 cells were provided by J.C. Marine [10] and were maintained in DMEM medium supplemented with 10% FBS, 1% P/S. The YUMM1.7 murine melanoma primary line (41) was grown in OPTIMEM medium with 10% FBS, 1% P/S. Lung (A549), breast (MDA-MB-231) and pancreatic (PANC1) cancer cells were grown in DMEM medium supplemented with 7% FBS, 1% P/S. For live imaging and red nuclear labeling 1205Lu cells were transduced with NucLight Red lentivirus reagent (Essen Bioscience) and selected with puromycin (1 μg/ml, Sigma Aldrich). Cells were maintained at 37°C in 5% CO2. pQCXIH-Myc-YAP-5SA was a gift from K. Guan (Addgene plasmid # 33093).

### Preparation of collagen gels

Soft and stiff collagen gels were made according to the protocol described in Park et al., 2020. Soft collagen gels have a stiffness <0.2 kPa and stiff collagen gels have a stiffness >4kPa. When not specified, cells were grown on stiff collagen gels.

For soft collagen gels, a solution containing 8 parts of rat type I collagen (Corning) concentrated to 2.5 mg/ml, 1 part of 10X PBS and 1 part of NaOH (0.1N) was used to obtain a basic pH favoring gel formation. This solution is then spread evenly with a sterile cone tip on the bottom of a well or culture diameter. The gel is then incubated for 1h at 37°C to allow it to polymerize. For stiff collagen gels, the rat type I collagen stock is diluted in 1X PBS to a final concentration of 50µg/ml. The solution is then deposited in the well or culture diameter and incubated for 1h at 37°C to allow it to polymerize. Cells are then detached using 1X collagenase I (Gibco) diluted in PBS according to the supplier’s specifications. The cells are then incubated for 20 minutes at 37°C to allow the collagenase I to digest the gel. A cell centrifugation is then performed to recover the cells in the pellet.

### RNAi studies

For transient RNAi transfection, cells were seeded on a culture dish and then transfected 24h later with 50nM siRNA targeting USP9X, YAP, β-TRCP or luciferase (non-targeting control) diluted in serum-free medium containing 5μM RNAi MAX (Thermo Fisher Scientific). Cells were then incubated at 37°C and 5% CO2 for the indicated time post transfection.

For the generation of cells stably depleted for USP9X, different shUSP9X sequences or a non-targeting shRNA sequence (directed against the Renilla luciferase mRNA) were cloned into a pSGEP Luc-PuroR lentiviral vector. Lentiviral particles were produced as follow. HEK293FT cells were seeded (500,000 cells in a T75 flask) and then transfected 72h later with 5µg of VSV-G plasmid, 10µg of pCMV 8.91 plasmid and 10µg of shUSP9X or shRenilla plasmid (control) diluted in optiMEM medium containing 60 µL of jetPEI DNA transfection reagent (Polyplus transfection). 1205Lu cells were seeded at 400,000 in a T75 flask. The medium containing the viral particles produced by HEK293FT was collected, centrifuged at 3,000 rpm for 5 minutes, filtered and put in contact with the 1205Lu cells 24 hours after seeding. Stably depleted cells were then selected with 1µg/ml puromycin.

### Immunoblot analysis

Cells were seeded on collagen gels. After pharmacological treatment or transfection, cells were lysed at 4°C in stop buffer (50mM TrisHCl, 150mM NaCl, 0.25% sodium deoxycholate, 1% NP-40, 1mM EDTA) supplemented with protease and phosphatase inhibitors (Pierce). Lysates were immersed in liquid nitrogen and centrifuged for 15 min at 13,300 rpm at 4°C. After colorimetric assay (DC protein assay-Biorad), 30 μg of total proteins are taken up in buffer (25mM Tris (pH 6.8), 2% SDS, 5% glycerol, 1% β-mercaptoethanol, 0.01% bromophenol blue), heated to 95°C 5min, plated on electrophoresis gels, separated, and transferred to nitrocellulose membrane (Millipore). Membranes were saturated 1h at RT in saturation buffer (5% BSA, 10mM Tris-HCl (pH7.5), 500mM NaCl) and incubated with primary antibodies at 4°C ON. The next day, membranes were rinsed in wash buffer (10mM Tris-HCl (pH7.5), 500mM NaCl, 0.1% Tween 20), incubated 1h at RT with the peroxidase-coupled secondary antibodies. After further washing, membranes were revealed by chemiluminescence (ECL, Millipore). Antibodies and inhibitors are listed in Supplementary Table 2.

### Affinity purification of poly-ubiquitinated proteins

Cell lysates were incubated with agarose-beads for 2h at 4°C and centrifuged at maximum speed to recover the supernatant. Pre-cleared supernatants were mixed with 5 µg of GST-TUBE (Tandem Ubiquitin Binding Entities, Life Sensors) fusion proteins with glutathione-agarose beads for 2h at 4°C pull-down ubiquitinated proteins. Samples were then centrifuged for 2 min at maximum speed. After 4 washes, pellets were then resuspended in 1X Laemli and analyzed according according to above immunoblot analysis.

### DUB TRAP assay and Immunoprecipitation

DUB TRAP assays were performed as described before (44). Briefly, cells were lysed at 4°C in buffer containing 50mM Tris (pH 7.4), 5mM MgCl2, 250Mm sucrose, 1mM DTT, 2mM ATP and 1mM PSMF. The lysates were immersed in liquid nitrogen and centrifuged 15min at 13,300 rpm at 4°C. Lysates were then incubated for 1h at 37°C with 0.5mM HA-Ub-VS (Boston Biochem), which label active DUBs by covalently binding with their enzymatic site. After this step, samples were processed according to above immunoblot analysis. Alternatively, lysates containing HA-Ub tagged DUBs were immunoprecipitated with anti-HA agarose beads (Sigma-Aldrich) for 16h at 4°C. After extensive washes in lysis buffer, beads were resuspended in Laemli buffer, boiled at 95°C for 5min and subjected to immunoblot analysis.

### Mass spectrometry analysis

For mass spectrometry analysis, anti-HA immunoprecipitated samples were loaded on NuPAGE™ 4–12% Bis–tris acrylamide gels according to the manufacturer’s instructions (Invitrogen, Life Technologies). Running of samples was stopped as soon as proteins stacked as a single band and following imperial blue staining (Life Technologies), the upper part of the gel containing the proteins was cut and processed for classical in gel digestion (washes, thiols reduction with 10 mM DTT and cystein alkylation with 55 mM iodoacetamide. Each band was further digested as previously described with trypsin and analyzed by liquid chromatography (LC)-tandem MS (MS/MS) using a Q Exactive Plus Hybrid Quadrupole-Orbitrap online with a nanoLC Ultimate 3000 chromatography system (Thermo Fisher Scientific™, San Jose, CA). For each biological sample (3 per condition), 4 microliters corresponding to 20 % of digested sample were injected in triplicate on the system. After pre-concentration and washing of the sample on a Acclaim PepMap 100 column (C18, 2 cm × 100 μm i.d. 100 A pore size, 5 μm particle size), peptides were separated on a LC EASY-Spray column (C18, 50 cm × 75 μm i.d., 100 A, 2 µm, 100A particle size) at a flow rate of 300 nL/min with a two steps linear gradient (2-22% acetonitrile/H20; 0.1 % formic acid for 100 min and 22-32% acetonitrile/H20; 0.1 % formic acid for 20 min). For peptides ionization in the EASYSpray source, spray voltage was set at 1.9 kV and the capillary temperature at 250 °C. All samples were measured in a data dependent acquisition mode. Each run was preceded by a blank MS run to monitor system background. The peptide masses were measured in a survey full scan (scan range 375-1500 m/z, with 70 K FWHM resolution at m/z=400, target AGC value of 3.00×10^6^ and maximum injection time of 100 ms). Following the high-resolution full scan in the Orbitrap, the 10 most intense data-dependent precursor ions were successively fragmented in HCD cell and measured in Orbitrap (normalized collision energy of 25 %, activation time of 10 ms, target AGC value of 1.00×10^5^, intensity threshold 1.00×10^4^ maximum injection time 100 ms, isolation window 2 m/z, 17.5 K FWHM resolution, scan range 200 to 2000 m/z). Dynamic exclusion was implemented with a repeat count of 1 and exclusion duration of 20 s. Data were processed according to the data processing protocol (Supplementary Methods).

### Proliferation assay

Real time analysis of cell proliferation was assessed using an automated videomicroscope (Incucyte Zoom system) according to the manufacturer’s instructions (Essen BioScience) and as described before by counting the nuclei of NucLight Red infected cells (36). Cells were seeded in triplicate in complete medium (15 x 10^3^ cells/well in 12-well plates) on stiff collagen substrate and treated with the indicated drugs. Phase contrast and red immunofluorescent images were taken every 6 h over a 5-day period (4 images per well at x 4 magnification). Cell proliferation was quantified by counting the number of fluorescent nuclei over time and growth curves were generated using the GraphPad prism 8 software. Results were normalized to time 0.

### Migration, invasion and *in vitro* scratch assays

Serum-stimulated chemotaxis and invasion were analysed using modified Boyden chambers (8 µm pores, Corning) as described before (63). For invasion assays, the upper side of the filter was coated with 0.5mg/ml matrigel (Corning, NY, USA). Migrated cells were stained with crystal violet and counted (four fields randomly per well). 3D Spheroid Invasion assays were performed as follow. A 30µl drop containing 3,000 cells was placed on the inside of the lid and incubated for 3 days using the hanging droplet method. Once the spheroid is formed, it was embedded in a mixture containing 2.5 mg/ml collagen, MEM 10X medium (Gibco), 1M HEPES and 0.3N NAOH to maintain pH 7. Invasion was monitored and then imaged at 24 h using a light microscope (Zeiss). The invasion area was quantified by Image J software. For *in vitro* scratch assays, cells were grown approximately 80% confluence in 24-well plates. Following treatment, wounds were generated using a sterile 200 μl pipette tip across each well and the coverage of the wound in each well was monitored by automated real-time imaging (Incucyte Zoom system).

### Time lapse analysis of cell locomotion

Cells were seeded on stiff or soft collagen gels in a 12-well plate (35,000 cells per well) and their nuclei were labeled with Hoescht. Cells were maintained at 37°C with 5% CO2 for 24 hours under a Leica TIRF videomicroscope and images were taken every 10min intervals for 24h. Cell movement was followed with the “ Trackmate “ plugin of the ImageJ software.

### Real-time quantitative PCR

Cells were seeded on stiff or soft collagen gels. After pharmacological treatment or transfection, total RNAs were extracted using the Nucleospin RNA plus kit according to the supplier’s recommendations (Macherey-Nagel). The amount of extracted RNA was measured using a Nanodrop (Thermo scientific). RNA was reverse transcribed using the High-Capacity cDNA reverse transcription kit (Applied Biosystems). Real-time amplification of amplicons corresponding to different target genes was followed on the Step One thermocycler (Applied Biosystems) by incorporation of SYBR Green (Applied Biosystems). Melting curve analysis was used to verify the specificity of the generated amplicons. The relative amount of each target was determined based on the expression of housekeeping genes using the ΔΔCt method. Heatmaps depicting fold changes of gene expression were prepared using MeV software.

### Immunofluorescence and microscopy

Cells were cultured in a 12-well plate on a collagen-coated 15mm glass coverslip (35,000 cells/condition). Cells were rinsed in PBS and fixed with 3% paraformaldehyde for 20 minutes. After washing, cells were incubated for 10 minutes with a buffer containing PBS + 0.3% Triton X100 for cell permeabilization and incubated in PBS, 0.1% Triton, 5% goat serum (Dako) for 1h, which allows saturation of non-specific sites. Cells were then labelled with primary antibodies in in PBS, 0.1% Triton. F-actin was stained with Alexa Fluor 488 phalloidin (1:100; Thermo Fisher Scientific) and nuclei were stained with DAPI. Following incubation with Alexa Fluor-conjugated secondary antibodies, coverslips were mounted in ProLong antifade mounting reagent (Thermo Fisher Scientific). Images were captured using a wide field microscope (Leica DM5500B). Quantification of the proportion of nuclear or cytoplasmic YAP was performed by comparing the nuclear fluorescence intensity to that of the cytoplasm using ImageJ software. Quantification of cell areas and YAP intensity were also performed with ImageJ software.

### In vivo experimentations, immunostaining and collagen imaging

All experiments requiring mice were performed in compliance with the institutional animal care and local animal ethics committees (CIEPAL-Azur agreement NCE/2018-509). For experimental lung metastasis assays, 5-week-old female nude mice (Janvier) were intravenously injected with 1205Lu Luc+ cells (1×10^6^) that were transduced with non-targeting shRNA lentivirus (shCTRL) or USP9X targeting shRNA lentivirus (shUSP9X) expressing a luciferase reporter gene. Images were recorded using a photon imager (Biospace lab) on mice injected intraperitoneally with 50mg/kg D-luciferin (PerkinElmer). Lung metastasis was monitored and quantified using BLI (62,63).

For tumor tracking experiments, 5-week-old C57BL/6 mice were injected with 5×10^5^ YUMM1.7 cells in 100*µL of PBS. Tumors were measured with caliper and treatments were started when the tumors reached a volume equivalent to 100mm3, after randomization of mice into control and test groups. Vemurafenib (30 mg/kg p.o.), Trametinib (0.3 mg/kg p.o.), USP9X inhibitor G9 (7.5 mg/kg i.p) or the indicated combinations were administered every 3 days in vehicle (90% corn oil, 10% DMSO). Control mice were treated with vehicle only. After animal sacrifice, tumors were dissected, weighed and snap-frozen in liquid nitrogen for RNA or protein extraction and immunofluorescence analysis (embedded in OCT from Tissue-Tek). For collagen imaging, tumor samples were fixed in formalin, embedded in paraffin and stained with picrosirius red using standard protocols. Tumor sections were analyzed using polarized light microscopy as previously described (35). For second harmonic generation (SHG) imaging, 7µM sections were deparaffinized and the collagen network was observed using a Zeiss 780NLO microscope (Carl Zeiss Microscopy) with Mai Tai HP DeepSee (Newport Corporation).

### Analysis of gene expression from public databases

USP9X interactome was generated using the STRING tool (https://string-db.org/network/9606.ENSP00000316357). Publicly available gene expression data sets of human melanoma cell lines were used to analyze *USP9X* and *YAP1* levels (GSE3189). Normalized data were analyzed using GraphPad Prism software. Pearson correlation between *USP9X* levels and *YAP1* transcriptional gene signature (CORDENONSI YAP conserved signature) was assessed using the skin melanoma TCGA database retrieved using cBioPortal (cbioportal.org). Survival data from the skin melanoma TCGA database were retrieved using ProteinAtlas (https://www.proteinatlas.org/).

### Statistical analysis

Unless otherwise stated, all experiments were repeated at least 3 times and representative data/images are shown. Statistical analysis was performed using using GraphPad Prism 8 software. The unpaired two-interval Mann-Whitney test was used for statistical comparisons between two groups when they do not follow a parametric distribution. The Student’s T test was used for samples following a parametric distribution. A Log-rank (Mantel-Cox) test was applied to Kaplan-Meier survival curves. Data represent biological replicates (n) and are depicted as mean values ± SEM as indicated in the figure legends.

## Data availability

The mass spectrometry proteomics data have been deposited on the ProteomeXchange Consortium of proteomics resource (http://www.proteomexchange.org) with the dataset identifier PXD054010.

## Acknowledgments

We thank J.C Marine for the short-term cultured melanoma cells, M. Irondelle from the C3M imaging facility, members of the C3M animal facility and C. Matthews for the SHG microscopy analysis (IBV imaging platform, UCA). This work was supported by funds from Inserm, Société Française de Dermatologie, Fondation ARC pour la recherche sur le cancer and Ligue Nationale Contre le Cancer. The financial contribution of the Conseil général 06, Canceropôle Provence Alpes Côte d’Azur and Région Provence Alpes Côte d’Azur to the C3M is also acknowledged. The Marseille Proteomic facility (MaP; http://map.univmed.fr/) is supported by IBiSA (Infrastructures Biologie Santé et Agronomie), Canceropôle PACA, Région PACA and Institut Paoli-Calmettes. P.B. was a recipient of a doctoral fellowship from Fondation ARC pour la recherche sur le cancer.

## Author contributions

M.D. and P.B. designed the study. P.B. performed the experiments and analyzed the data with the help of A.C., C.A.G., W.M., M.O. S.D., M.L. and O.B. S.A. performed the mass spectrometry analysis. M.K. provided lentiviral shRNA constructs. M.D. and P.B. wrote the original draft. M.D. supervised the study and edited the final version of the manuscript with the help of S.T.-D.

