## Supplemental files for "USP9X is a mechanosensitive deubiquitinase that controls tumor cell invasiveness and drug response through YAP stabilization"

**Supplementary Material and Methods**

**Supplementary Methods**

**Cell contraction assay**

Cells (5 x $10^4$) were embedded in 100 µL of collagen I/matrigel and seeded in a glass bottom 96-well plate (MatTek). Once the gel was set (1h at 37 C), cells were maintained in DMEM 10% FBS with or without indicated drugs. Gel contraction was monitored at day 3. The gel area was measured using ImageJ software and the percentage of contraction was calculated as described in (Girard 2020).

**Mass spectrometry data processing protocol**

Relative intensity-based label-free quantification (LFQ) intensity was processed using the MaxLFQ algorithm from the freely available MaxQuant computational proteomics platform, version 1.6.3.4. The acquired raw LC Orbitrap MS data were first processed using the integrated Andromeda search engine. Spectra were searched against the human database subset of the SwissProt database (date 20210424, 20395 entries). This database was supplemented with a set of 245 frequently observed contaminants. The following parameters were used for searches: (i) trypsin allowing cleavage before proline; (ii) two missed cleavages were allowed; (ii) monoisotopic precursor tolerance of 20 ppm in the first search used for recalibration, followed by 4.5 ppm for the main search and 0.5 Da for fragment ions from MS/MS ; (iii) cysteine carbamidomethylation (+57.02146) as a fixed modification and methionine oxidation (+15.99491) and N-terminal acetylation (+42.0106) as variable modifications; (iv) a maximum of five modifications per peptide allowed; and (v) minimum peptide length was 7 amino acids and a maximum mass of 4,600 Da. The match between runs option was enabled to transfer identifications across different LC-MS/MS replicates based on their masses and retention time within a match time window of 0.7 min and using an alignment time window of 20 min. The quantification was performed using a minimum ratio count of 1 (unique+razor) and the second peptide option to allow identification of two co-fragmented co-eluting peptides with similar masses. The false discovery rate (FDR) at the peptide and protein levels were set to 1% and determined by earching a reverse database. For protein grouping, all proteins that cannot be distinguished based on their identified peptides were assembled into a single entry according to the MaxQuant rules. The statistical analysis was done with Perseus program (version 1.6.15) from the MaxQuant environment (www.maxquant.org). The LFQ normalised intensities were uploaded from the ProteinGroups.txt file. Proteins marked as contaminant, reverse hits, and “only identified by site” were discarded. Quantifiable proteins were defined as those detected in 70% of samples in at least one condition among all conditions. Protein intensities were base 2 logarithmized to obtain a normal distribution. Missing values were replaced using data imputation by randomly selecting from a normal distribution centered on the lower edge of the intensity values that simulates signals of low abundant proteins using default parameters (a downshift of 1.8 standard deviation and a width of 0.3 of the original distribution). In this way, imputation of missing values in the controls allows statistical comparison of protein abundances that are present only in the positive Immunoprecipitated samples and absent in negative control. To determine whether a given detected protein was specifically differential a two-sample t-test were done using permutation based FDR-controlled at 0.01 and employing 250 permutations. The p value was adjusted using a scaling factor s0 with a value of 1. The differential proteomics analysis was carried out on identified proteins after removal of proteins only identified with modified peptides, peptides shared with other proteins, proteins from contaminant database and protein which are present above 70% in at least one condition.

**Supplementary Figure legends**

**Supplementary Figure 1.** Effect of collagen matrix stiffness on DUB activity on A549 (lung cancer cells), MDA-MB-231 (breast cancer cells) and PANC1 (pancreas cancer cells) cells cultivated for 72h on soft versus stiff collagen matrices. Lysates of the indicated cells were incubated at 37°C with the HA-Ub-VS probe and analyzed by anti-HA. Arrowhead indicates the position of USP9X. ERK2, loading control.

**Supplementary Figure 2.** Biochemical analysis of USP9X and YAP nuclear localization in 1205lu cells grown on soft or stiff collagen matrix. Lamin, nuclear marker. RHO GDI, cytoplasmic marker.

**Supplementary Figure 3. USP9X controls YAP expression and activity in multiple solid cancers. A.** Western blot analysis of USP9X and YAP expression on A549, MDA-MB-231 and PANC1 cells transfected for 72h with the siUSP9X#1, the siYAP#1 or a control siRNA and cultivated on stiff collagen matrices. ERK2, loading control. **B.** Time-lapse microscopy analysis of A549, MDA-MB-231 and PANC1 cells displacement during 24h after siUSP9X or siControl transfection. Upper panel, displacement path of 5 representative cells for each condition. Lower panel, quantification of cell displacement (n>40). ****p<0.0001, ***p<0.001, *p<0.05, Kruskal-Wallis analysis.

**Supplementary Figure 4. Targeting USP9X impairs MAPKi-induced cell contractile activity and potentiates the action of targeted therapies *in vitro*. A.** Collagen contraction assays of 1205Lu cells pretreated for 72 hours with BRAFi (1µM vemurafenib) in the presence or not of the indicated doses of USP9X inhibitor G9. Gel contraction was monitored after 3 days. Bar graph represents the mean ± SD (n=5). *p<0.05, ns, non-significant, unpaired two-tailed Mann–Whitney's test. **B.** 1205lu melanoma cells were seeded on stiff collagen matrices, treated with the combination of BRAFi (1µM vemurafenib) and MEKi (0.1 µM trametinib) in the presence or not of USP9X inhibitor (0.5µM G9) and proliferation was monitored using real time imaging (Incucyte Zoom System). Curves show the evolution of confluency through time relative to T0. Data are the mean ± SD from a minimum of 10 fields per conditions. ****p<0.0001, ns, non-significant, two-way ANOVA analysis. **C.** Western blot analysis of USP9X, YAP, cleaved Caspase 3, p27 and phosphorylated ERK on lysates from cells treated for 72h with the combination of BRAFi (1µM vemurafenib) and MEKi (0.1 µM trametinib) in the presence or not of the indicated dose of USP9X inhibitor (G9) on stiff collagen matrices. ERK2, loading control.

**Supplementary Table 2**

**Antibodies**

| **Antibody** | **Source** | **Reference** |
|---|---|---|
| aSMA | Abcam | ab9694 |
| B-TRCP | Santa Cruz | sc-33213 |
| YAP | Cell Signaling Technology | 14074S |
| Fibronectin | Sigma | F3648 |
| HSP90 | Santa Cruz | sc-13119 |
| YAP/TAZ | Cell Signaling Technology | 84185 |
| USP9X | Novus biological | NB100-61591 Novus |
| ERK2 | Santa Cruz | sc-154 |
| FAP | Santa Cruz | Sc-65398 |
| SLUG | Santa Cruz | Sc-10436 |
| HA | Cell Signaling Technology | 3724S |
| CYR61 | Santa Cruz | sc-374129 |
| SERP1 | Santa Cruz | sc-5297 |
| THBS1 | Cell Signalling Technology | 376795 |
| ANLN | | GTX107742 |
| Phospho-ERK | Cell Signalling Technology | |
| ACTIN | | |

**Pharmacological inhibitors**

| Compound | Source | Reference |
|---|---|---|
| G9 | Selleckchem | S6877 |
| Cycloheximide | Sigma | 01810-G |
| Bortezomib | Selleckchem | S1013 |
| Vemurafenib | AbMole | M1761 |
| Trametinib | AbMole | M1759 |
| Blebbistatine | Selleckchem | S7099 |

| Rank | Protein names | Gene names | Student's T-test Difference Stiff_Soft |
|---|---|---|---|
| **DUBs enriched on Stiff** | | | |
| **1** | OTU domain-containing protein 5 | OTUD5 | 2,722674635 |
| **2** | Ubiquitin carboxyl-terminal hydrolase 35 | USP35 | 2,228630834 |
| **3** | Ubiquitin carboxyl-terminal hydrolase 38 | USP38 | 1,870641973 |
| **4** | Ubiquitin carboxyl-terminal hydrolase CYLD | CYLD | 1,509820117 |
| **5** | Ubiquitin thioesterase OTUB2 | OTUB2 | 1,501484685 |
| **6** | **Probable ubiquitin carboxyl-terminal hydrolase FAF-Y** | **USP9Y** | **1,480303261** |
| **7** | Ubiquitin carboxyl-terminal hydrolase 32 | USP32 | 1,410126421 |
| **8** | Ubiquitin carboxyl-terminal hydrolase isozyme L5 | UCHL5 | 1,37678406 |
| **9** | Ubiquitin carboxyl-terminal hydrolase 16 | USP16 | 1,371511062 |
| **10** | OTU domain-containing protein 6B | OTUD6B | 1,343094534 |
| **11** | Ubiquitin carboxyl-terminal hydrolase 8 | USP8 | 1,326086071 |
| **12** | Ubiquitin carboxyl-terminal hydrolase 24 | USP24 | 1,319959508 |
| **13** | Ubiquitin carboxyl-terminal hydrolase 15 | USP15 | 1,317871915 |
| **14** | Ubiquitin carboxyl-terminal hydrolase 47 | USP47 | 1,286602789 |
| **15** | Ubiquitin carboxyl-terminal hydrolase 10 | USP10 | 1,26393395 |
| **16** | Ubiquitin carboxyl-terminal hydrolase 14 | USP14 | 1,248148017 |
| **17** | Ubiquitin carboxyl-terminal hydrolase 4 | USP4 | 1,222306861 |
| **18** | Ubiquitin carboxyl-terminal hydrolase isozyme L3 | UCHL3 | 1,209354692 |
| **19** | Ubiquitin carboxyl-terminal hydrolase 22 | USP22 | 1,18237967 |
| **20** | Ubiquitin carboxyl-terminal hydrolase 5 | USP5 | 1,17518017 |
| **21** | Ubiquitin carboxyl-terminal hydrolase 25 | USP25 | 1,171805408 |
| **22** | Ubiquitin carboxyl-terminal hydrolase 28 | USP28 | 1,152943002 |
| **23** | Ubiquitin carboxyl-terminal hydrolase 19 | USP19 | 1,118481212 |
| **24** | OTU domain-containing protein 7B | OTUD7B | 1,078283283 |
| **25** | **Probable ubiquitin carboxyl-terminal hydrolase FAF-X** | **USP9X** | **1,048494524** |
| **26** | Ubiquitin thioesterase OTU1 | YOD1 | 1,030632151 |
| **27** | Ubiquitin carboxyl-terminal hydrolase 40 | USP40 | 1,028741625 |
| **28** | OTU domain-containing protein 1 | OTUD1 | 1,018856843 |
| **DUBs enriched on Soft** | | | |
| **1** | Lys-63-specific deubiquitinase BRCC36 | BRCC3 | -2,659423722 |
| **2** | mor necrosis factor alpha-induced protein 3;A20p50;A20p | TNFAIP3 | -2,04895197 |
| **3** | U4/U6,U5 tri-snRNP-associated protein 2 | USP39 | -1,899883986 |
| **4** | COP9 signalosome complex subunit 6 | COPS6 | -1,207294517 |
| **5** | COP9 signalosome complex subunit 5 | COPS5 | -1,02998315 |

**Supplementary Table 1. List of deubiquitinases (DUBs) controlled by extracellular matrix stiffness in melanoma cells.** Lysates from 1205Lu cells grown on soft or stiff collagen matrices were incubated with an activity-based HA-tagged ubiquitin. Lysates were immunoprecipitated with anti-HA agarose beads. Immunoprecipitates were then analysed by LC-MS/MS analysis. Red background, DUBs found enriched when cells are plated on stiff collagen matrix. Green background, DUBs found enriched when cells are plated on soft collagen matrix.

### Supplementary Figure 1

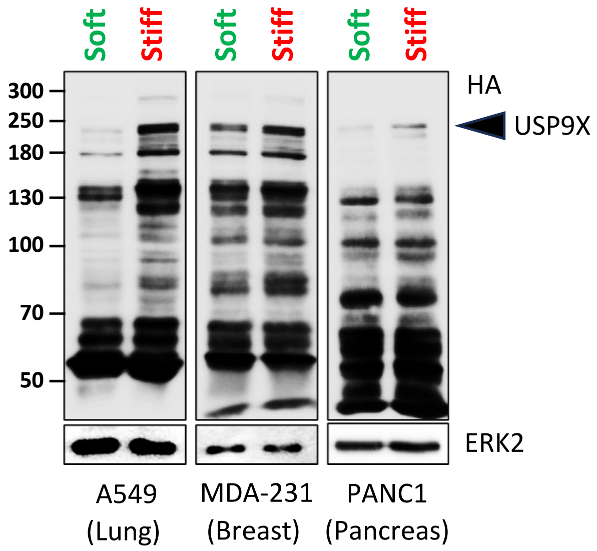

### Supplementary Figure 2

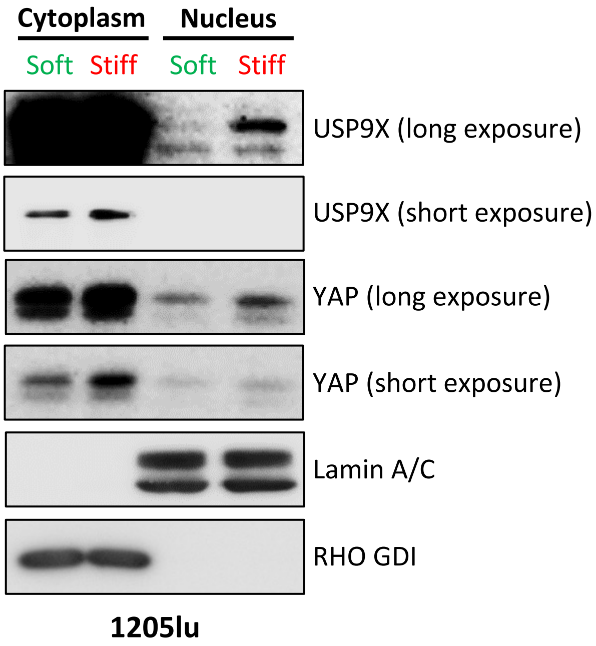

### Supplementary Figure 3

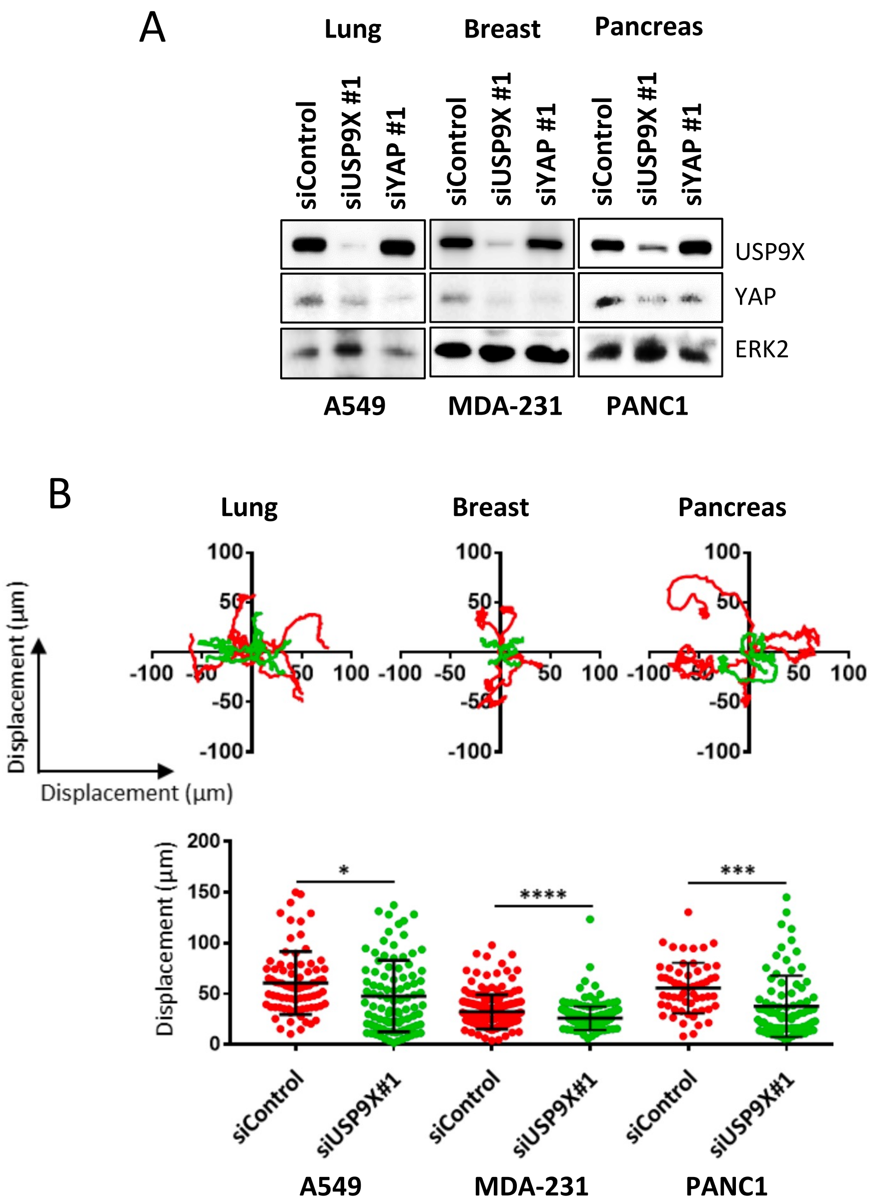

### Supplementary Figure 4

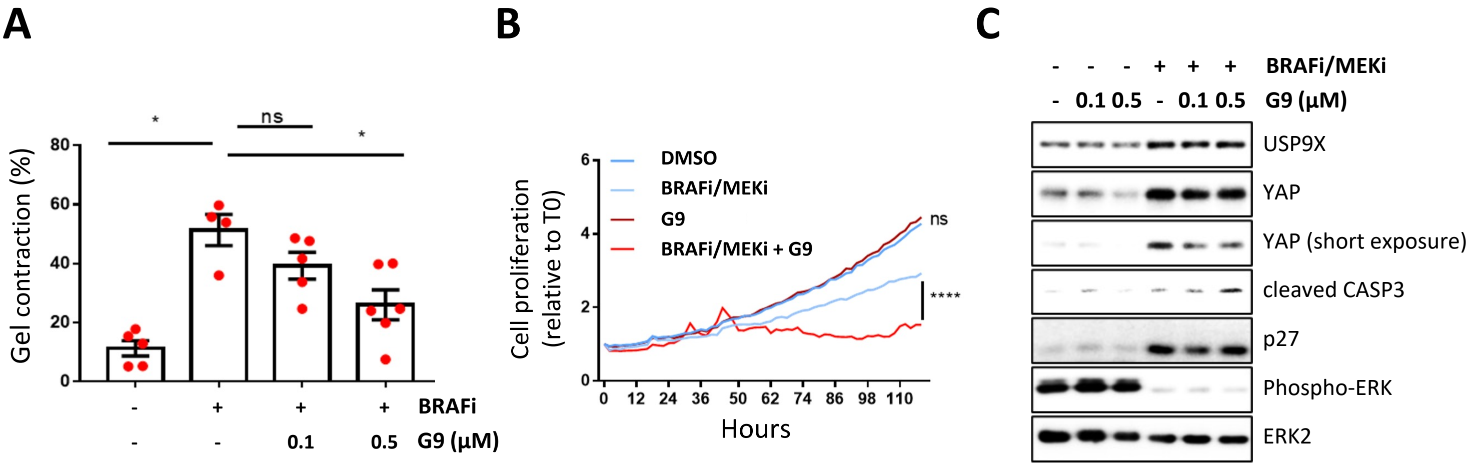
